# Metabolic Reprogramming from Glycolysis to Fatty Acid Uptake and beta-Oxidation in Platinum-Resistant Cancer Cells

**DOI:** 10.1101/2021.05.17.444564

**Authors:** Junjie Li, Yuying Tan, Guangyuan Zhao, Kai-Chih Huang, Horacio Cardenas, Daniela Matei, Ji-Xin Cheng

## Abstract

Increased aerobic glycolysis is widely considered as a hallmark of cancer. Yet, cancer cell metabolic reprograming during development of therapeutic resistance is under-studied. Here, through high-throughput stimulated Raman scattering imaging and single cell analysis, we found that cisplatin-resistant cells exhibit increased uptake of exogenous fatty acids, accompanied with decreased glucose uptake and de novo lipogenesis, indicating a reprogramming from glucose and glycolysis dependent to fatty acid uptake and beta-oxidation dependent anabolic and energy metabolism. A metabolic index incorporating measurements of glucose derived anabolism and fatty acid uptake correlates linearly to the level of resistance to cisplatin in ovarian cancer cell lines and in primary cells isolated from ovarian cancer patients. Mechanistically, the increased fatty acid uptake facilitates cancer cell survival under cisplatin-induced oxidative stress by enhancing energy production through beta-oxidation. Consequently, blocking fatty acid beta-oxidation by a small molecule inhibitor in combination with cisplatin or carboplatin synergistically suppressed ovarian cancer proliferation in vitro and growth of patient-derived xenograft in vivo. Collectively, these findings support a new way for rapid detection of cisplatin-resistance at single cell level and a new strategy for treatment of cisplatin-resistant tumors.

## INTRODUCTION

Metabolic reprogramming in cancer cells has been recognized since the discovery of the Warburg effect in 1920s ^1, 2^. Increased aerobic glycolysis is now widely considered as a hallmark of many cancers and clinically exploited as a target for cancer therapy and a cancer biomarker for diagnosis ^3^. In the past decade, numerous studies have investigated the heterogeneity and complexity of cancer metabolism beyond the Warburg effect ^4^. Metabolic reprogramming allows cancer cells to adapt to intrinsic or extrinsic cues from the microenvironment through plasticity and high flexibility in nutrient acquisition and utilization ^5^. Particular attention has been paid to metabolic alterations associated with critical steps of cancer progression, such as metastasis initiation, circulation and colonization ^5–7^. Metabolic reprogramming in cancer stem cells identified potential vulnerabilities for cancer stem cells targeting therapy ^8, 9^. Cancer cells also rewire their metabolic dependencies within a specific microenvironment niche by interacting with stroma cells ^10^ or with the surrounding adipocytes ^11, 12^. Further, alterations in nutrient utilization under metabolic stress conditions have recently been reported ^13–15^. Despite these recent advances, the understanding of cancer cell metabolism remains far from complete. One of the less studied areas is cancer metabolic reprogramming associated with resistance to therapy.

Therapeutic resistance remains one of the biggest challenges facing cancer treatment. Resistance to chemotherapy or molecularly targeted therapies is a major cause of tumor relapse and death ^16^. Emerging studies support an association between metabolic reprogramming and cancer drug resistance ^17, 18^. Several studies have linked the Warburg effect to resistance to radiation ^19^ and lactate production was shown to promote resistance to chemotherapy in cervical cancer ^20^. Altered lipid metabolism has also been implicated in acquisition of drug resistance ^21^. Increased de novo lipogenesis mediated by FASN facilitated gemcitabine resistance in pancreatic cancer ^22^ while cancer associated adipose tissue promoted resistance to antiangiogenic interventions by supplying fatty acids to cancer cells in regions where the glucose demand was insufficient^23^. Additionally, lipid droplet production mediated by lysophosphatidylcholine acyltransferase 2 promoted resistance of colorectal cancer cells to 5-fluorouracil and oxaliplatin ^24^. Recently, it has been proposed that drug tolerant cells adopt a state of diapause similar to suspended embryonic development to survive chemotherapy toxic insults, in which cell proliferation and metabolic processes are suppressed ^25^. These studies support that metabolic reprograming underlie development of drug resistance and point to potential metabolic vulnerabilities of resistant cancer cells, which remain under utilized.

Platinum-based drugs, including cisplatin, carboplatin and oxaliplatin, represent one class of the most widely used chemotherapy drugs ^26^. Resistance to platinum is a barrier to effective treatment in multiple cancers, including ovarian, testicular, bladder, head and neck, non-small-cell lung cancer and others ^27^. Understanding the metabolic reprograming underlying platinum resistant cancer cells is critical for development of effective treatment strategies. Yet, precisely profiling such metabolic reprogramming using conventional technology is difficult, because within a large cell population, only a small portion of cells is drug resistant or tolerant. In this study, by taking advantage of a hyperspectral stimulated Raman scattering (SRS) imaging platform, we depict the metabolic profile of platinum resistant cancer cells at the single cell level.

SRS microscopy is a recently developed label-free chemical imaging technique that detects the intrinsic chemical bond vibrations ^28–31^. The value of SRS microscopy was demonstrated in identifying cholesteryl ester accumulation as a signature associated with multiple aggressive cancers ^32, 33^, discovering increased lipid desaturation in ovarian cancer stem cells ^8^, and tracing metabolic flux by isotope labeling ^34, 35^. More recently, large-area hyperspectral SRS microscopy and high-throughput single cell analysis revealed lipid-rich protrusion in cancer cells under stress ^36^. Raman spectro-microscopy based single cell metabolomics unveiled an important role of lipid unsaturation in aggressive melanoma ^37^.

Here, through large-area hyperspectral SRS imaging and subsequent single-cell analysis, we identified a stable metabolic switch from glucose and glycolysis dependent to fatty acid uptake and beta-oxidation dependent anabolic and energy metabolism in cisplatin-resistant ovarian cancer cells. By coupling metabolic flux through isotope labeling and SRS-based molecular imaging, we found that cisplatin-resistant cells display increased uptake of exogenous fatty acids, accompanied with decreased glucose uptake and *de novo* lipogenesis. By incorporating SRS imaging-based measurements of glucose derived anabolism and fatty acid uptake, we introduce a metabolic index defined as the ratio of fatty acid uptake versus glucose incorporation. The metabolic index was found linearly correlated to the level of resistance to cisplatin in ovarian cancer cell lines and in primary cells, demonstrating a potential of using SRS imaging for rapid detection of cisplatin-resistance *ex vivo*. Mechanistically, the cisplatin-resistant cells display higher fatty acid oxidation rate, which supplies additional energy and promotes cancer cell survival under cisplatin-induced oxidative stress. Blocking beta-oxidation by a small molecule inhibitor or genetic perturbation in combination with cisplatin or carboplatin treatment synergistically suppressed ovarian cancer proliferation *in vitro* and growth of a patient-derived xenograft model *in vivo*. In addition, acute treatment with cisplatin induced a transient metabolic shift towards higher fatty acid uptake in lung, breast, and pancreatic cancers. Together, these results promise new treatment options for patients with cisplatin-resistant tumors by targeting the fatty acid beta-oxidation pathway.

## RESULTS

### High-throughput SRS imaging of lipid metabolism in single living ovarian cancer cells

To identify the altered lipid metabolism in cisplatin-resistant cells, we established a high-throughput single-cell analysis approach, which couples large-area hyperspectral SRS scanning of 200-500 cells per group with spectral phasor segmentation and CellProfiler analysis. As shown in **Supplementary Fig. 1a**, we firstly acquire a stack of large-area hyperspectral SRS images, containing hundreds of individual cells in each field-of-view. An SRS spectrum is extracted at each pixel from the image stack. Then, the hyperspectral SRS images are segmented through a spectral phasor algorithm to generate maps of intracellular compartments corresponding to nuclei and lipids (mostly in lipid droplets) based on the spectrum similarity ^38^. Next, the nuclei map is inputted into CellProfiler ^38^ to guide the identification of the edges of each individual cell from the raw whole cell image. After individual cells are outlined, the lipid map is mapped back to the corresponding cells. Finally, quantitative characterization of lipids in terms of integrated intensity, mean intensity, area, and lipid droplet size in each individual cell is performed.

To explore the lipid metabolic signature of cisplatin-resistant ovarian cancer cells, we generated isogenic pairs of cisplatin-resistant cells from three ovarian cancer cell lines, including SKOV3, OVCAR5, and COV362, through long-time exposure and recovery after cisplatin treatment at IC50 concentration ^39^. Resistance to cisplatin in these cell lines was validated by repeat assays measuring cisplatin dose response. All the resistant cell lines exhibited 2~3-fold increase of IC_50_ when compared to their parental counterpart cell lines (**Supplementary Fig. 1b-e**). In addition, we studied isogenic PEO1/PEO4 cell lines derived from the same patient, at the time of a platinum sensitive (PEO1) and platinum resistant-recurrence (PEO4) ^40^.

Taking advantage of the high-throughput imaging analysis platform, we analyzed the lipid metabolism in these 4 pairs of cisplatin-resistant and parental ovarian cancer cell lines. Comparing SRS images of sensitive PEO1 and cisplatin-resistant PEO4 cells, there tends to be an increase of lipid intensity in PEO4 cells but with large cell-to-cell variations (**Fig. 1a**). We quantitatively analyzed the integrated lipid intensity in individual cells and plotted them in histograms. Intriguingly, the histograms display two distinct subpopulations, lipid-poor and lipid-rich, in each cell line, implying a metabolic heterogeneity within the same group. While lipid-poor cells dominate in PEO1 cell line, PEO4 cells show a dramatic increase in lipid-rich subpopulation and a decrease in lipid-poor subpopulation (**Fig. 1b**). Single-cell analysis reveals an even more obvious increase of lipid-rich subpopulation and decrease of lipid-poor subpopulations in SKOV3-cisR cells, compared to SKOV3 (**Fig. 1c and 1d**). Additionally, we observed similar lipid content change in the other two pairs of cell lines, including OVCAR5 versus OVCAR5-cisR (**Supplementary Fig. 1f**), and COV362 versus COV362-cisR (**Supplementary Fig. 1g**), implying that cisplatin-resistant cells harbor higher level of lipid accumulation. Furthermore, after acute treatment with cisplatin, a significant increase of lipid-rich subpopulation and decrease of lipid-poor subpopulation are found in SKOV3 cells (**Fig. 1e**), but no obvious change of lipid distribution pattern in SKOV3-cisR cells was detected (**Fig. 1f**), supporting that lipid-rich cells are more resistant to cisplatin treatment. These data collectively suggest that higher level of lipid content is a metabolic feature of cisplatin-resistant ovarian cancer cells.

**Fig. 1:**
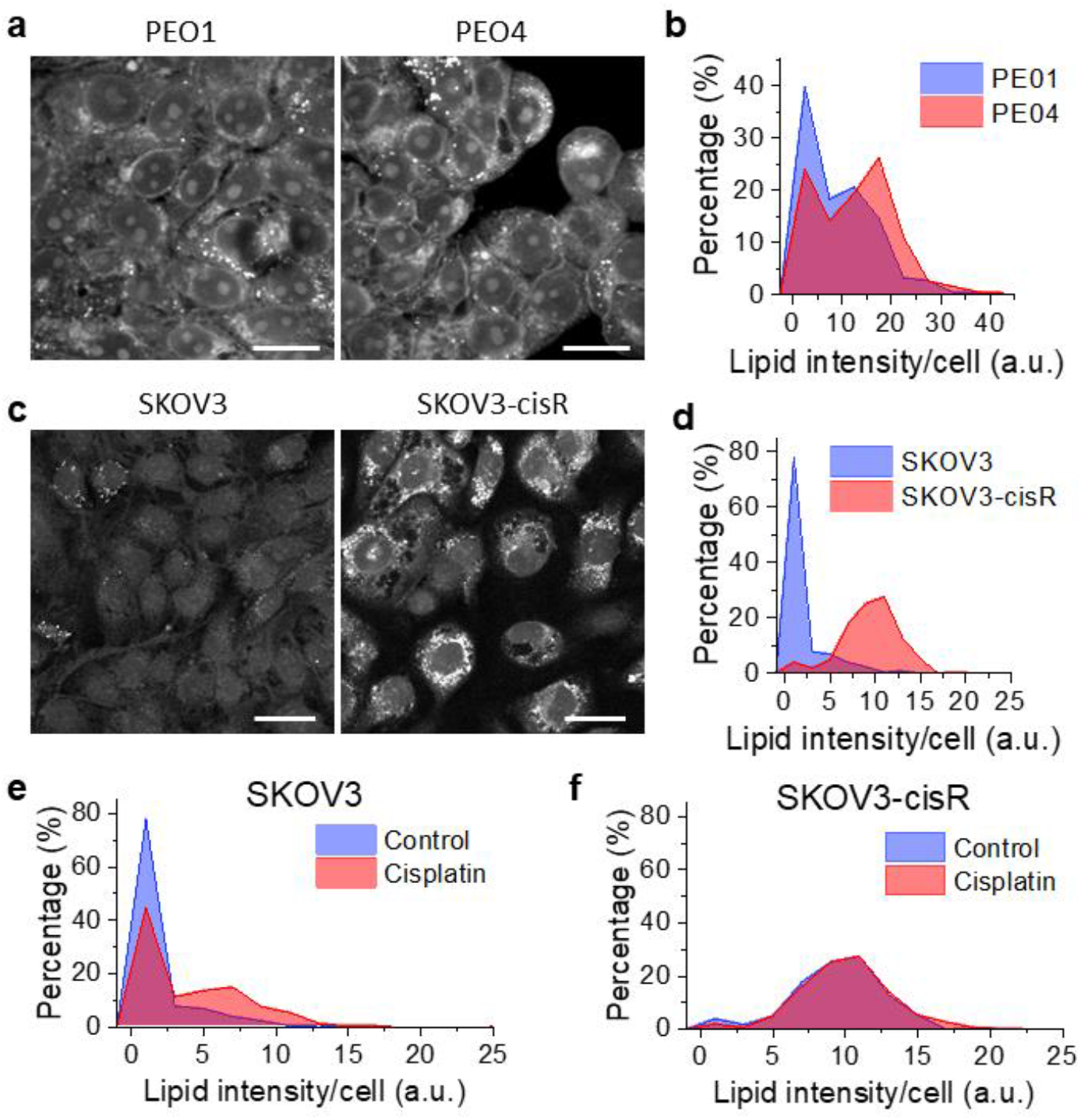
High-throughput imaging of lipid metabolism in isogenic pairs of cisplatin-sensitive and -resistant ovarian cancer cells. **(a)** Representative large-area SRS images of parental PEO1 and cisplatin-resistant PEO4 cells. Scale bar: 20 µm. **(b)** Histograms of integrated cellular lipid intensity in PEO1 and PEO4 cells generated through high-throughput single-cell analysis. **(c)** Representative large-area SRS images of parental SKOV3 and cisplatin-resistant SKOV3-cisR cells. Scale bar: 100 µm. **(d)** Histograms of integrated cellular lipid intensity in SKOV3 and SKOV3-cisR cells. **(e)** Histograms of integrated cellular lipid intensity in SKOV3 cells treated with or without cisplatin. **(f)** Histograms of integrated cellular lipid intensity in SKOV3-cisR cells treated with or without cisplatin.

### Increased fatty acid uptake but not de novo lipogenesis contributes to high-level lipid content in cisplatin-resistant ovarian cancer cells

To identify the source of increased lipid content in cisplatin-resistant ovarian cancer cells, we examined the contribution of *de novo* lipogenesis and of fatty acid uptake to lipid accumulation, respectively. By a stable isotope probing method ^35^, we examined the level of lipogenesis by feeding the cells with deuterium labeled glucose-d_7_. Newly synthesized macromolecules (mostly lipids) were imaged by hyperspectral SRS microscopy at Raman shift from 2050 cm^−1^ to 2350 cm-1, covering the vibrational frequencies of C-D bonds. SRS images show weaker C-D signal in cisplatin-resistant PEO4 cells than the signal in parental PEO1 cells (**Fig. 2a**). Quantitative analysis confirms significant reduction of both signal intensity and relative area fraction in PEO4 cells when compared to PEO1 cells (**Fig. 2b**), indicating a decrease in glucose derived anabolism and *de novo* lipogenesis in cisplatin-resistant cells. Using a similar approach, we examined the fatty acid uptake level by hyperspectral SRS imaging at C-D vibrational frequencies in cells fed with deuterium labeled palmitic acid-d_31_ (PA-d_31_). In contrast to glucose-d_7_ fed cells, C-D signal in PA-d_31_ fed PEO4 cells is stronger than PEO1 cells (**Fig. 2c**). Quantitative analysis revealed significant increase of both signal intensity and relative area fraction (**Fig. 2d**). In addition to saturated fatty acid (PA-d_31_), we further tested the uptake of an unsaturated fatty acid, oleic acid-d_34_ (OA-d_34_). Uptake of OA-d_34_ was significantly increased in PEO4 cells comparing to PEO1 cells (**Fig. 2e and 2f**). These results suggest the increased uptake of fatty acid is not specific to certain type of fatty acid, but rather reflects an upregulation of fatty acid uptake pathway.

**Fig. 2:**
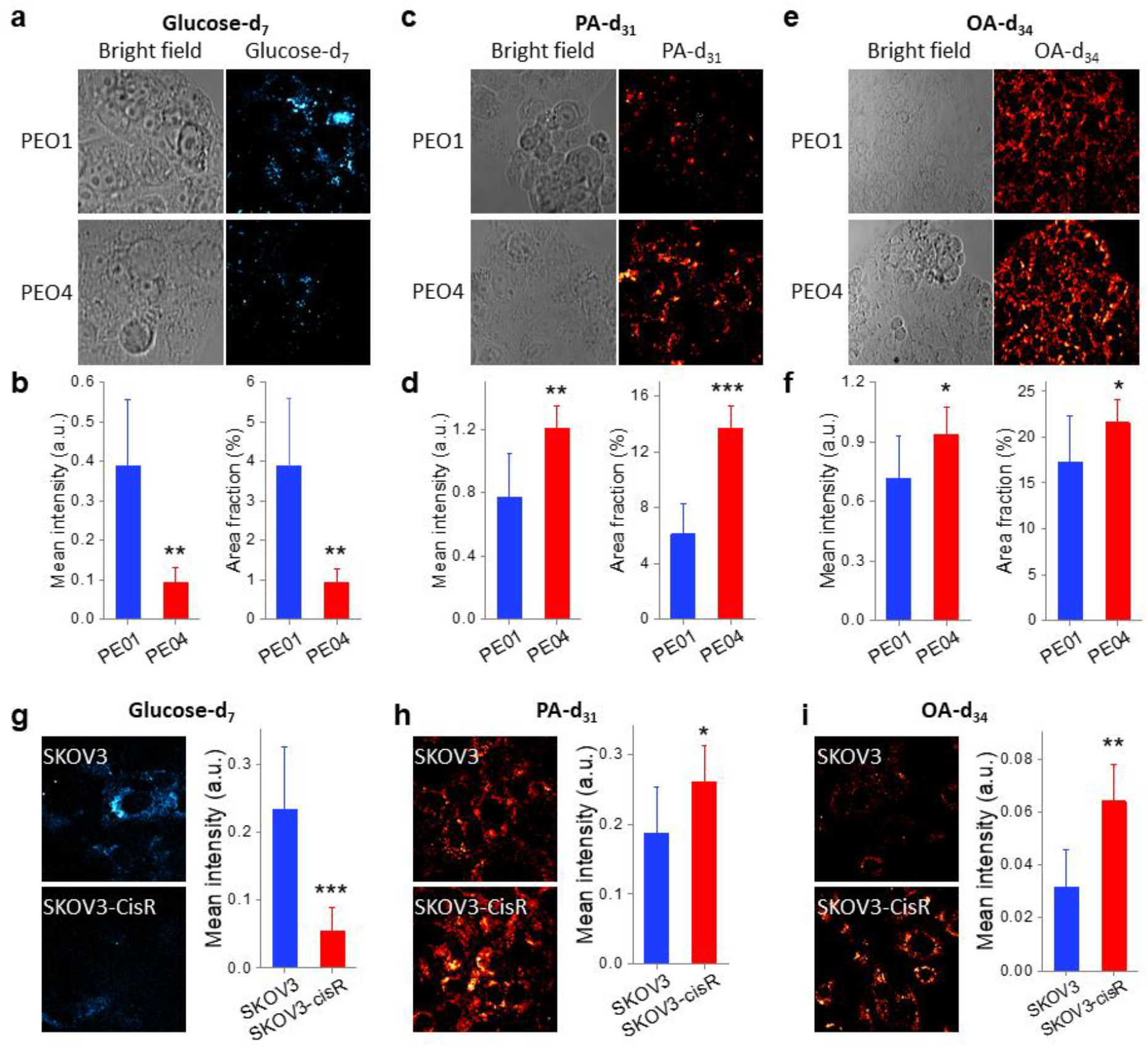
Increased fatty acid uptake, not de novo lipogenesis, is the major contributor to lipid accumulation in cisplatin-resistant ovarian cancer cells. **(a)** Representative bright field and SRS images of PEO1 and PEO4 cells fed with glucose-d_7_ for 3 days. **(b)** Quantitative analysis of SRS signal of C-D bonds in glucose-d_7_ fed PEO1 and PEO4 cells by mean intensity and area fraction. **(c)** Representative bright field and SRS images of PEO1 and PEO4 cells fed with PA-d_31_ for 6 h. **(d)** Quantitative analysis of SRS signal of C-D bonds in PA-d_31_ fed PEO1 and PEO4 cells by mean intensity and area fraction. **(e)** Representative bright field and SRS images of PEO1 and PEO4 cells fed with OA-d_34_ for 6 h. **(f)** Quantitative analysis of SRS signal of C-D bonds in OA-d_34_ fed PEO1 and PEO4 cells by mean intensity and area fraction. **(g)** Representative SRS images of SKOV3 and SKOV3-cisR cells fed with glucose-d_7_ for 3 days and quantitative analysis of SRS signal of C-D bonds by mean intensity. **(h)** Representative SRS images of SKOV3 and SKOV3-cisR cells fed with PA-d_31_ for 6 h and quantitative analysis of SRS signal of C-D bonds by mean intensity. **(i)** Representative SRS images of SKOV3 and SKOV3-cisR cells fed with OA-d_34_ for 6 h and quantitative analysis of SRS signal of C-D bonds by mean intensity. The results in all the column graphs are shown as means + SD, n = 8-10. * *P* < 0.05, ** *P* < 0.01, and *** *P* < 0.001.

To verify if the observed phenomenon is cell type specific, we repeated the measurements in SKOV3 and SKOV3-cisR cells. Consistently, SRS images and quantitative analysis showed a significant decrease in glucose-d_7_ derived C-D signal in SKOV3-cisR cells (**Fig. 2g**) and increase in PA-d_31_ signal (**Fig. 2h**) and OA-d34 signal in SKOV3-cisR cells (**Fig. 2i**), when compared to parental SKOV3 cells. Additionally, we observed the same trend in the other two pairs of cell lines, OVCAR5 versus OVCAR5-cisR (**Supplementary Fig. 2a**) and COV362 versus COV362-cisR (**Supplementary Fig. 2b**). These data collectively suggest a metabolic switch from glucose derived anabolism to fatty acid uptake in cisplatin-resistant ovarian cancer cells.

### Metabolic index as a predictor of cisplatin resistance

Having shown decreased glucose-derived anabolism and increased fatty acid uptake in cisplatin-resistant ovarian cancer cells, we continued exploring whether this metabolic feature can be used for differentiation of cisplatin-resistant from cisplatin-sensitive cancer cells. To quantitatively characterize the resistance to cisplatin, we calculated IC_50_ dose of cisplatin in various cell lines and their C-D intensities (presented as area fraction) from glucose-d_7_, PA-d_31_, or OA-d_34_. (**Table S1**). Interestingly, glucose-d_7_ derived C-D intensity was found negatively correlated to IC_50_ to cisplatin (**Fig. 3a**), while PA-d_31_ intensity was positively correlated to IC_50_ to cisplatin (**Fig. 3b**). To integrate two measurements into one, we used the ratio of PA-d_31_/(PA-d_31_ + Glucose-d_7_), giving a dimensionless number ranging from 0 to 1. We defined this ratio as the “metabolic index”. The index linearly correlated to the IC_50_ to cisplatin (**Fig. 3c**), suggesting its potential to detect and quantitatively determine resistance to cisplatin in cancer cells.

**Fig. 3:**
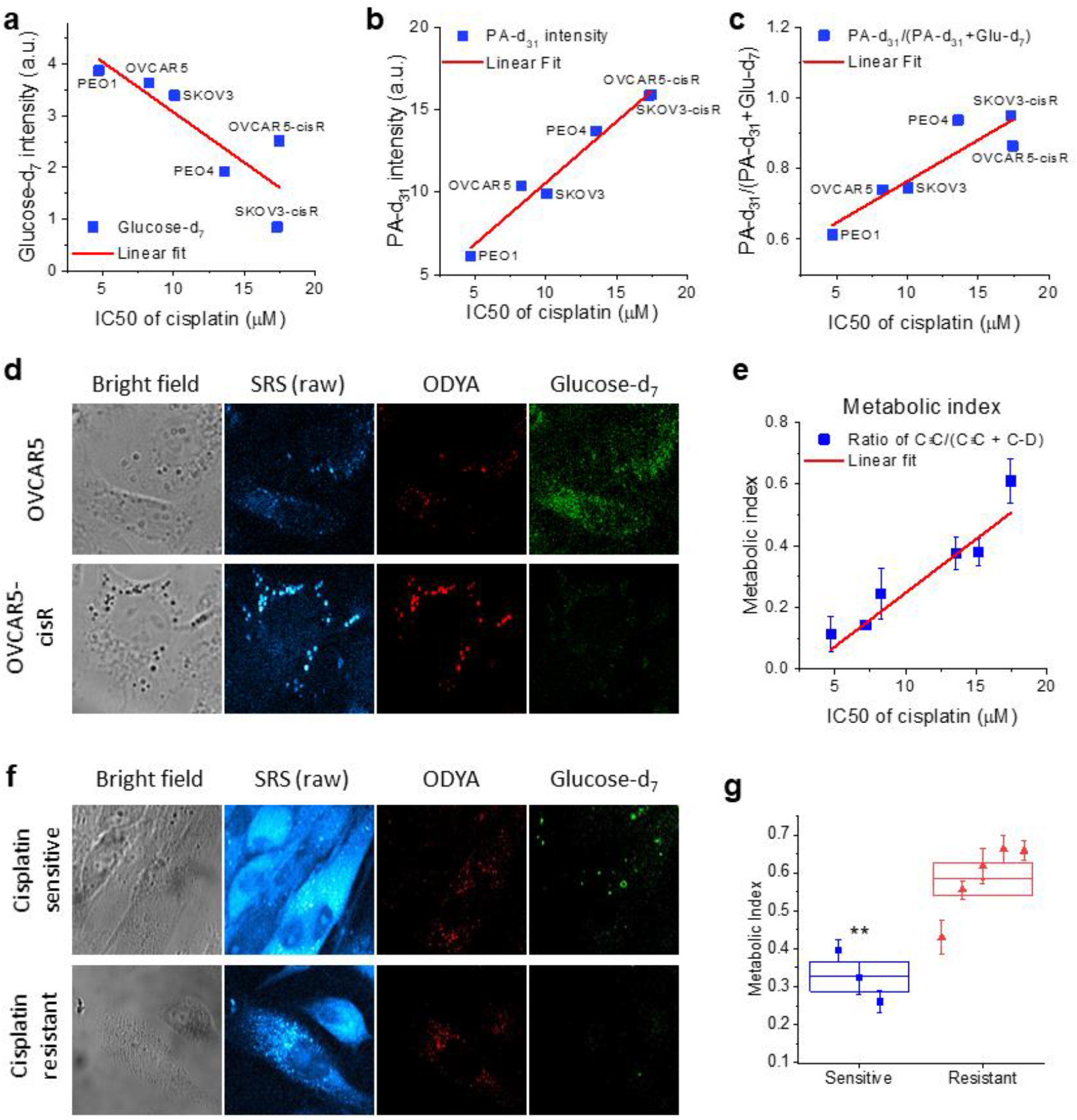
Metabolic index by integrating glucose derived lipogenesis and fatty acid uptake directly correlates with cisplatin resistance. **(a)** A linear regression of glucose-d_7_ intensity to IC_50_s of cisplatin in various ovarian cancer cell lines. R^2^ = 0.6596. **(b)** A linear regression of PA-d_31_ intensity to IC_50_s of cisplatin in various ovarian cancer cell lines. R^2^ = 0.9611. **(c)** A linear regression of the ratio of PA-d_31_/(PA-d_31_ + Glucose-d_7_) to IC_50_s of cisplatin in various ovarian cancer cell lines. R^2^ = 0.7808. **(d)** Representative bright field images, raw SRS images, and processed SRS images of ODYA and glucose-d_7_ in OVCAR5 and -cisR cells. **(e)** A linear regression of the metabolic index, as defined by the ratio of C≡C/(C≡C + C-D) to IC_50_s of cisplatin in various ovarian cancer cell lines. R^2^ = 0.9235. **(f)** Representative bright field images, raw SRS images, and processed SRS images of ODYA and glucose-d_7_ in primary ovarian cancer cells from cisplatin treatment resistant patients and from the cisplatin treatment sensitive patient.**(g)**Quantitative analysis of metabolic index (the ratio of C≡C/(C≡C + C-D) for primary ovarian cancer cells from cisplatin treatment resistant patients and from the cisplatin treatment sensitive patient. Each data point represents the average metabolic index of individual cancer cells from a patient and its error bar indicates the SEM; n=11-31. The boxes are shown as means ± SEM for each patient group. ** *P* < 0.01.

Understanding the value of this metabolic imaging method for detecting cisplatin resistance at the single cell level, we further developed a method that allowed us to simultaneously image both glucose derived anabolism and fatty acid uptake in the same cell. Instead of using deuterium labeled fatty acids to trace fatty acid uptake, we used a fatty acid analog, 17-octadecynoic acid (ODYA). ODYA has an endogenous C≡C at one end of the fatty acid chain, which produces a strong Raman peak around 2100 cm^−1^ (**Supplementary Fig. 3a**). The distinctive Raman spectrum of ODYA enables separation of C≡C labeled fatty acid (from fatty acid uptake) from C-D labeled macromolecules derived from glucose-d_7_ (**Supplementary Fig. 3a**). To test this, we performed hyperspectral SRS imaging in cells fed with both ODYA and glucose-d_7_. From images, we observed two signals with distinctive spectra, from C≡C labeled fatty acids and C-D derived from glucose-d_7_, respectively (**Supplementary Fig. 3b**).

Next, we applied this approach to image OVCAR5 and OVCAR5-cisR cells. Two components, C≡C and C-D, were segmented from the raw SRS images based on the spectral shasor algorithm ^38^. Consistently, we observed apparently stronger C≡C signal and weaker C-D signal in OVCAR5-cisR cells, when compared to OVCAR5 cells (**Fig. 3d**). Quantitative analysis confirms a significant increase of C≡C and decrease of C-D signal. The metabolic index, ratio of C≡C/(C≡C + C-D), is more significantly increased in OVCAR5-cisR cells (**Supplementary Fig. 3c**). Following this established protocol, we analyzed metabolic indices in the other three pairs of cell lines, including PEO1 and PEO4 (**Supplementary Fig. 3d and S3e**), SKOV3 and SKOV3-cisR (images not shown), and COV362 and COV362-cisR cells (images not shown). Consistently, a linear correlation was established between metabolic index and IC_50s_ of cisplatin in these cell lines (**Fig. 3e**).

To further validate the metabolic index as a predictor of platinum resistance in clinically relevant samples, we applied this method to primary ovarian cancer cells obtained from human patients with known status of resistance/response to platinum. As shown in **Fig. 3f**, in ovarian cancer cells isolated from a patient with platinum-sensitive disease, signal from ODYA was observed only in some of the cells, but the signal from glucose-d_7_ was relatively strong. In ovarian cancer cells isolated from cisplatin-resistant tumors, ODYA signal was more evenly distributed in the cells imaged, while signal from glucose-d_7_ was weaker. Quantitative analysis showed that the metabolic indices are higher in samples from five patients with resistant tumors, when compared to cancer cells from three patients with sensitive disease (**Fig. 3g**). While studies from a larger pool of patients are needed to fully explore the applicability of metabolic index measurement to detect platinum resistance, our pilot studies suggest the potential of metabolic imaging for predicting platinum resistance in tumor samples from patients.

### Fatty acid uptake contributes to cisplatin resistance

Knowing that cisplatin-resistant ovarian cancer cells uptake more fatty acids, we asked whether the fatty acid uptake is a cause or result of cisplatin resistance. Firstly, we tested whether modulating exogenous fatty acid availability affect endogenous lipid amount in cisplatin-resistant ovarian cancer cells. We cultured OVCAR5-cisR cells in lipid-deficient culture medium or normal medium supplemented with 1% lipid mixture for 24 h and then examined lipid amount by SRS microscopy. The results showed that the lipid-deficiency significantly reduced intra-cellular lipid amount while lipid supplementation increased the lipid amount (**Fig. 4a and 4b**). Similar phenomenon was observed in the SKOV3-cisR cell line (**Supplementary Fig. 4a and S4b**). These observations further support the forementioned conclusion that fatty acid uptake, instead of *de novo* lipogenesis, is the major source of the lipid accumulation in cisplatin-resistant ovarian cancer cells. Next, we examined whether modulating exogenous lipid availability impacts cancer cell’s resistance to cisplatin. We found that lipid-deficiency increased sensitivity to cisplatin, while lipid supplementation slightly decreased sensitivity to cisplatin in OVCAR5-cisR (**Fig. 4c**), PEO4 (**Supplementary Fig. 4c**), and SKOV3-cisR cells (**Supplementary Fig. 4d**). These data suggest that resistance to cisplatin can be alleviated by modulating exogenous fatty acid availability.

**Fig. 4:**
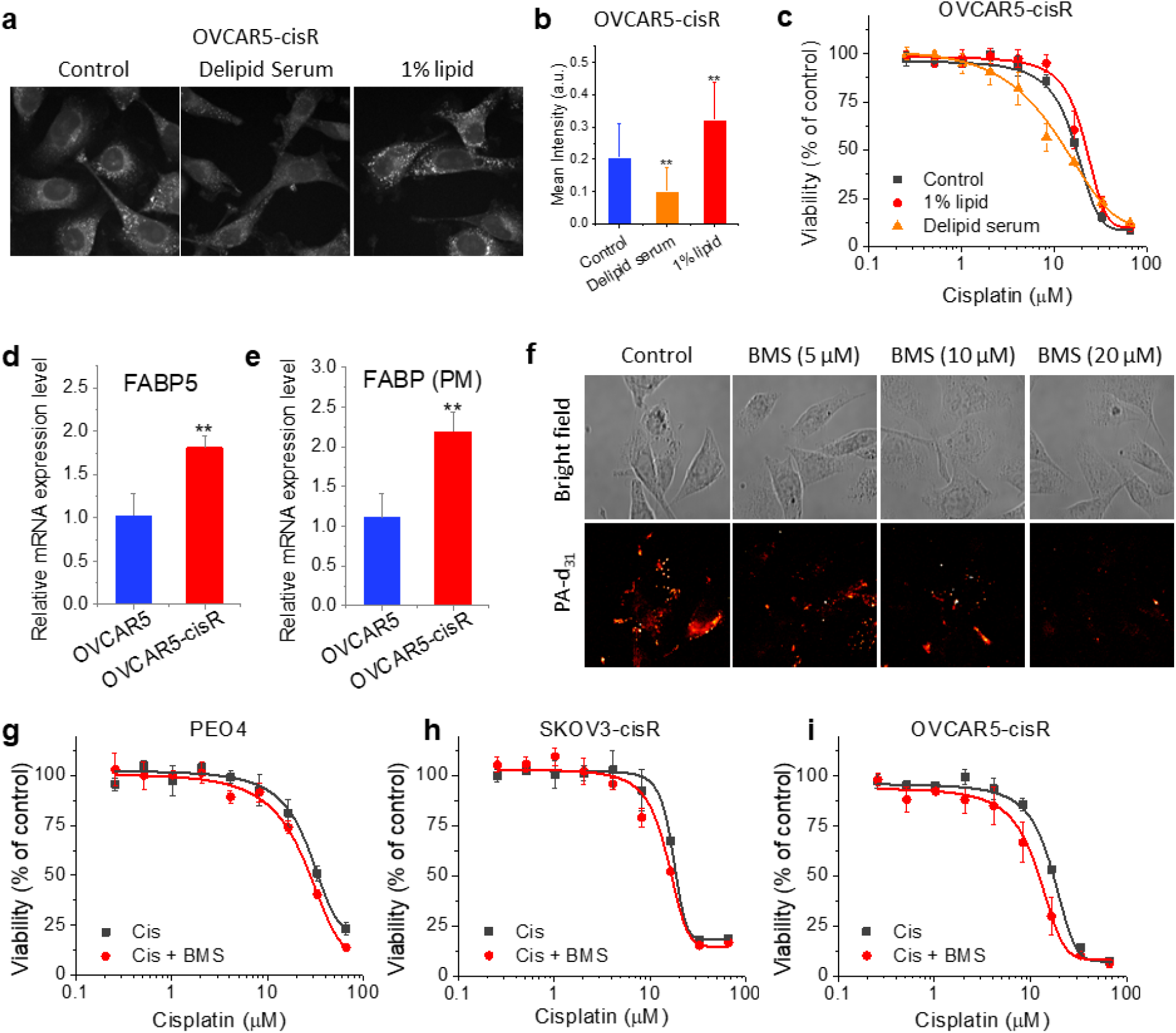
Fatty acid uptake directly contributes to cisplatin resistance. **(a)** SRS image of OVCAR5-cisR cell cultured with control serum (FBS), delipid serum or control serum supplemented with 1% lipid mixture for 24 hours. **(b)** Quantitative C-H signal from lipid droplet of OVCAR5-cisR cell cultured with control serum (FBS), delipid serum or control serum supplemented with 1% lipid for 24 hours. **(c)** Dose-response to cisplatin under culture environment with control, reduced (medium containing delipid serum or no serum) and increased (control serum supplemented with 1% lipid mixture) lipid content for OVCAR5-cisR cells. **(d-e)** Relative mRNA express level of FABP5 (d) and FABP(PM) (e) in OVCAR5 and -cisR cells. The results are shown as means + SD; n = 3. **(f)** Representative bright field and SRS images of SKOV3-cisR cell after FA transporter inhibitor BMS309403 (BMS) treatment at 5 μM, 10 μM or 20 μM for 24 hours with the incubation of 100 μM PA-d31 for 6 hours. **(g-i)** Dose-response to cisplatin with or without supplemental BMS treatment for PEO4 (g), SKOV3-cisR (h) and OVCAR5-cisR (i) cell. The results in all the does-response curves are shown as means ± SD; n =3. The results in all the column graphs are shown as means + SD, n = 8-10. * *P* < 0.05, ** *P* < 0.01, and *** *P* < 0.001.

Fatty acid uptake is a cellular process facilitated by multiple fatty acid transporters/carriers, including CD36, FATPs, and FABPs ^41, 42^. One of the proteins that have been reported to be upregulated in ovarian cancer is the fatty acid binding protein 4 (FABP4) ^12, 43^. We asked whether the increased fatty acid uptake in cisplatin-resistant ovarian cancer cells is regulated through upregulation of FABP4. We observed very low FABP4 mRNA levels in OVCAR5 and OVCAR5-cisR cells, suggesting FABP4 is not likely to play a major role in mediating the increased fatty acid uptake in cisplatin-resistant OVCAR5 cells. Next, we investigated the expression of a panel of other fatty acid uptake regulator genes, including CD36, FATP1-6, FABP5, and GOT2 (FABP PM) ^44^. FABP5 and FABP PM were found to be highly expressed in OVCAR5-cisR compared to parental cells, suggesting that they could be mediating fatty acid uptake in cisplatin-resistant cells (**Fig. 4d-e** and **Supplementary Fig. 4e**).

Next, we tested whether a potent inhibitor of FABP, BMS-309403, can suppress fatty acid uptake in cisplatin-resistant cells ^45^. Treatment with BMS-309403 significantly reduced PA-d_31_ uptake in SKOV3-cisR cells in a dose dependent manner (**Fig. 4f and S4f**). Suppression of fatty acid uptake by BMS-309403 was also observed in OVCAR5-cisR cells (**Supplementary Fig. 4g and S4h)**. Furthermore, inhibition of fatty acid uptake by BMS-309403 significantly reduced resistance to cisplatin in multiple resistant cell lines, including PEO4 (**Fig. 4i**), SKOV3-cisR (**Fig. 4j**), and OVCAR5-cisR cells (**Fig. 4k**), which supports a functional role of fatty acid uptake in the development of cisplatin resistance in ovarian cancer cells.

### Increased fatty acid oxidation rate contributes to cisplatin resistance

Considering that one major function of lipids is energy production through beta-oxidation, we next investigated whether fatty acid oxidation is increased in cisplatin resistant cancer cells. By measuring the oxygen consumption rate (OCR) in parental and resistant cancer cells, we found OVCAR5-cisR cells displayed much higher level of oxygen consumption than OVCAR5 cells (**Fig. 5a**). To test whether the increased oxidation rate arises from fatty acid oxidation, we used etomoxir, an inhibitor of CPT1, which transports fatty acid into mitochondria for beta-oxidation.

**Fig. 5:**
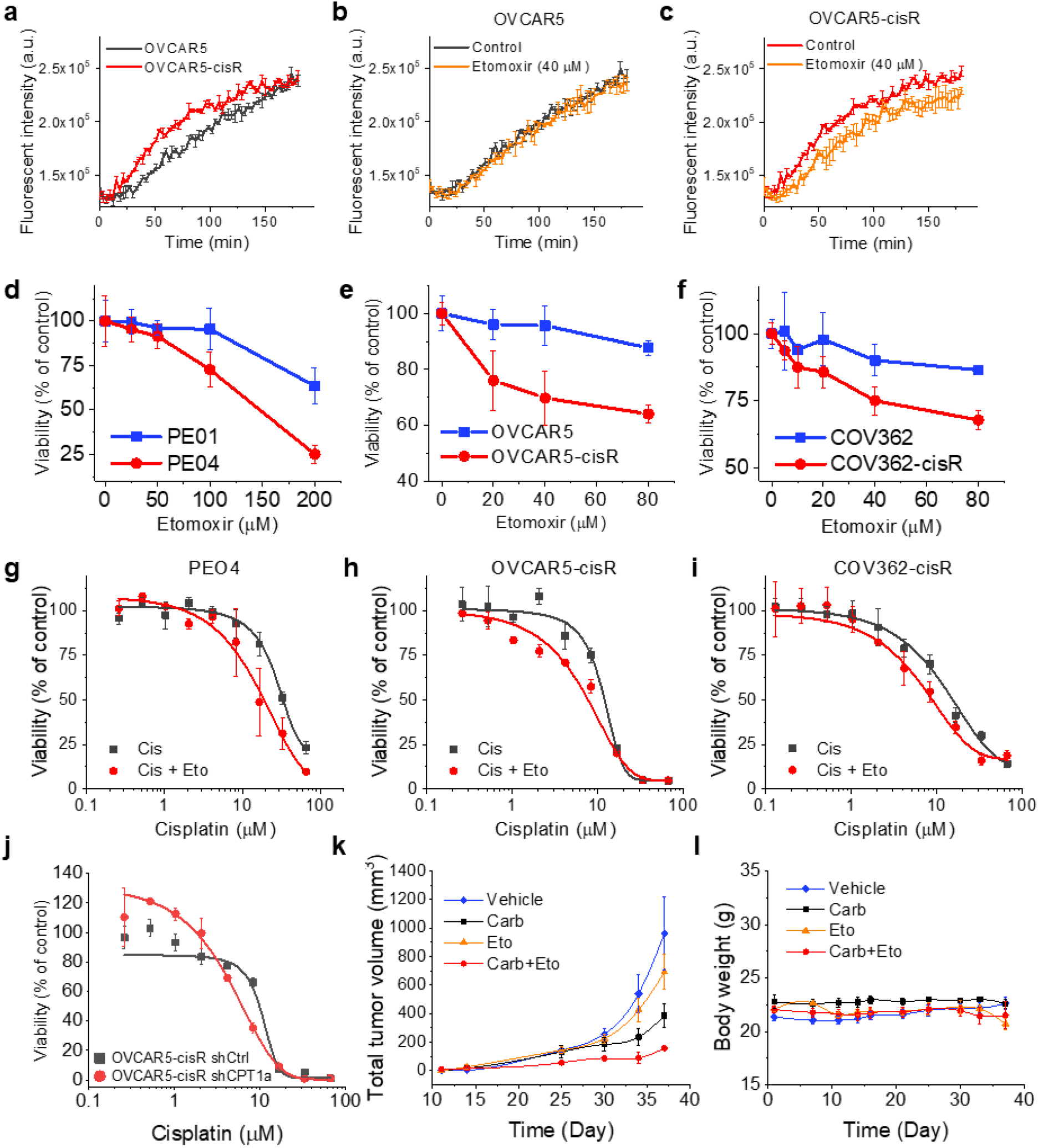
Fatty acid uptake contributes to cisplatin resistance by increasing fatty acid oxidation. **(a)** Oxygen consumption curves of OVCAR5-cisR and OVCAR5 over 3 hours **(b-c)** Oxygen consumption curves of OVCAR5-cisR (b) and OVCAR5 (c) with 40 μM etomoxir treatment over 3 hours. **(d-f)** Dose-response to etomoxir for cisplatin resistant cell lines and their parental cell lines including PEO1 and PEO4 (d), OVCAR5 and -cisR (e) and COV362 and -cisR (f). **(g-i)** Dose-response to cisplatin with or without supplemental etomoxir treatment at 40 μM for PEO4 (g), OVCAR5-cisR (h) and COV362 -cisR (i) cell. **(j)** Dose-response to cisplatin for OVCAR5-cisR shCtrl and shCPT1a cell. The data in all the does-response curves are shown as means ± SD; n = 3. **(k)** Total tumor volume growth curve from day 14 to 37 after tumor cell inoculation for vehicle, carboplatin, etomoxir and combinational treatment groups. **(l)** Mice body weight record since tumor inoculation for vehicle, carboplatin, etomoxir and combinational treatment groups. The data for PDX *in vivo* experiment are shown as means ± SEM; n = 3-6.

Etomoxir did not induce an obvious change in oxygen consumption in OVCAR5 cells (**Fig. 5b**), but significantly reduced oxygen consumption in OVCAR5-cisR cells (**Fig. 5c**). Quantitation of OCR confirmed a significant reduction of OCR after etomoxir treatment in OVCAR5-cisR cells, but not in OVCAR5 cells (**Supplementary Fig. 5a**). Additionally, Seahorse measurement of OCR in PEO1 and PEO4 cells supports a significant increase of fatty acid oxidation rate in PEO4 cells compared to PEO1 cells (**Supplementary Fig. 5b**). These data indicate that fatty acid oxidation significantly increases in cisplatin-resistant cells.

To test whether the increased fatty acid oxidation contributes to cisplatin resistance, we firstly tested the response to etomoxir in cisplatin-resistant cells and parental cells. We observed higher sensitivity to etomoxir treatment in cisplatin-resistant cell lines when compared to their parental cell lines, in paired cell lines including PEO1 and PEO4 (**Fig. 5d**), OVCAR5 and OVCAR5-cisR (**Fig. 5e**), and COV362 and COV362-cisR (**Fig. 5f**), indicating higher dependence on fatty acid oxidation in cisplatin-resistant cells. Next, we tested whether etomoxir treatment could reduce resistance to cisplatin. Our data show that the dose-response to cisplatin curves were significantly left shifted with etomoxir treatment in PEO4 (**Fig. 5g**), OVCA5-cisR (**Fig. 5h**), and COV362-cisR cells (**Fig. 5i**) and support potentially using etomoxir to re-sensitize resistant ovarian cancer cells to cisplatin. The observation was further confirmed by shRNA mediated knockdown of CPT1a in OVCAR5-cisR cells (**Supplementary Fig. 5c and S5d**). Knockdown of CPT1a in OVCAR5-cisR increased its vulnerability to cisplatin treatment compared to the control group (**Fig. 5j**). However, the CPT1a mRNA and protein levels were similar in OVCAR5 and OVCAR5-cisR cells (**Supplementary Fig. 5e and S5f)**, which implies the enhanced fatty acid oxidation in cisplatin-resistant ovarian cancer cell is likely the result of increased activation of CPT1a. This is consistent with our observation of decreased de novo lipogenesis in cisplatin-resistant ovarian cancer cells, which releases inhibition on CPT1 from malonyl-CoA, an intermediate of de novo lipogenesis ^46, 47^.

To test whether interruption of fatty acid oxidation could sensitize ovarian cancer to platinum, we used a patient-derived xenograft (PDX) model of ovarian cancer rendered platinum resistant through repeated exposure to carboplatin *in vivo* and described previously ^39^. To avoid toxicity induced by cisplatin, we substituted cisplatin with carboplatin, a second-generation agent. Carboplatin or etomoxir single-agent treatment induced a slight reduction of tumor growth, whereas the combination treatment caused dramatic suppression of tumor growth (**Fig. 5k**). Importantly, body weights remained stable in all groups, suggesting that the combination treatment was tolerable (**Fig. 5l**). SRS imaging of tissues extracted from the PDX model shows extensive lipid storage in cancer cells *in vivo* (**Supplementary Fig. 5g**). These data collectively support development of a combination of a platinum with a fatty acid oxidation inhibitor for platinum-resistant cancer treatment.

### Fatty acid oxidation facilitates cancer cell survival under cisplatin-induced oxidative stress

Cisplatin has been known to cause cytotoxicity by inducing oxidative stress, in addition to DNA adduct formation ^48–50^. Excess oxidative stress can inhibit glycolysis by inactivating key glycolytic enzymes, such as pyruvate kinase M2 (PKM2) and glyceraldehyde 3-phosphate dehydrogenase (GAPDH) ^15, 51^. Increased reactive oxidative species (ROS) oxidizes intracellular NADPH and thus suppresses *de novo* lipogenesis, as NADPH is one of the precursors for lipogenesis ^52^. Therefore, we hypothesize that fatty acid uptake and oxidation promote cancer cell survival under cisplatin-induced oxidative stress by replenishing free fatty acids and ATP, deficiency of which is caused by decreased *de novo* lipogenesis and glycolysis under oxidative stress. To test this hypothesis, we first checked the oxidative stress level by measuring intracellular ROS by using a fluorescent probe, 2’,7’-dichlorodihydrofluorescein diacetate (H2DCFDA). By confocal microscopy, we found that OVCAR5-cisR cells showed much stronger fluorescent signal than OVCAR5 cells (**Fig. 6a-b**). Similar trend of increased ROS in PEO4 cells compared to PEO1 cells was also observed (**Supplementary Fig. 6a and S6b**). Furthermore, we analyzed the change in ROS in OVCAR5 and OVCAR5-cisR cells treated with cisplatin and found that cisplatin treatment induced significant increase of ROS production in both cell lines (**Fig. 6c**).

**Fig. 6:**
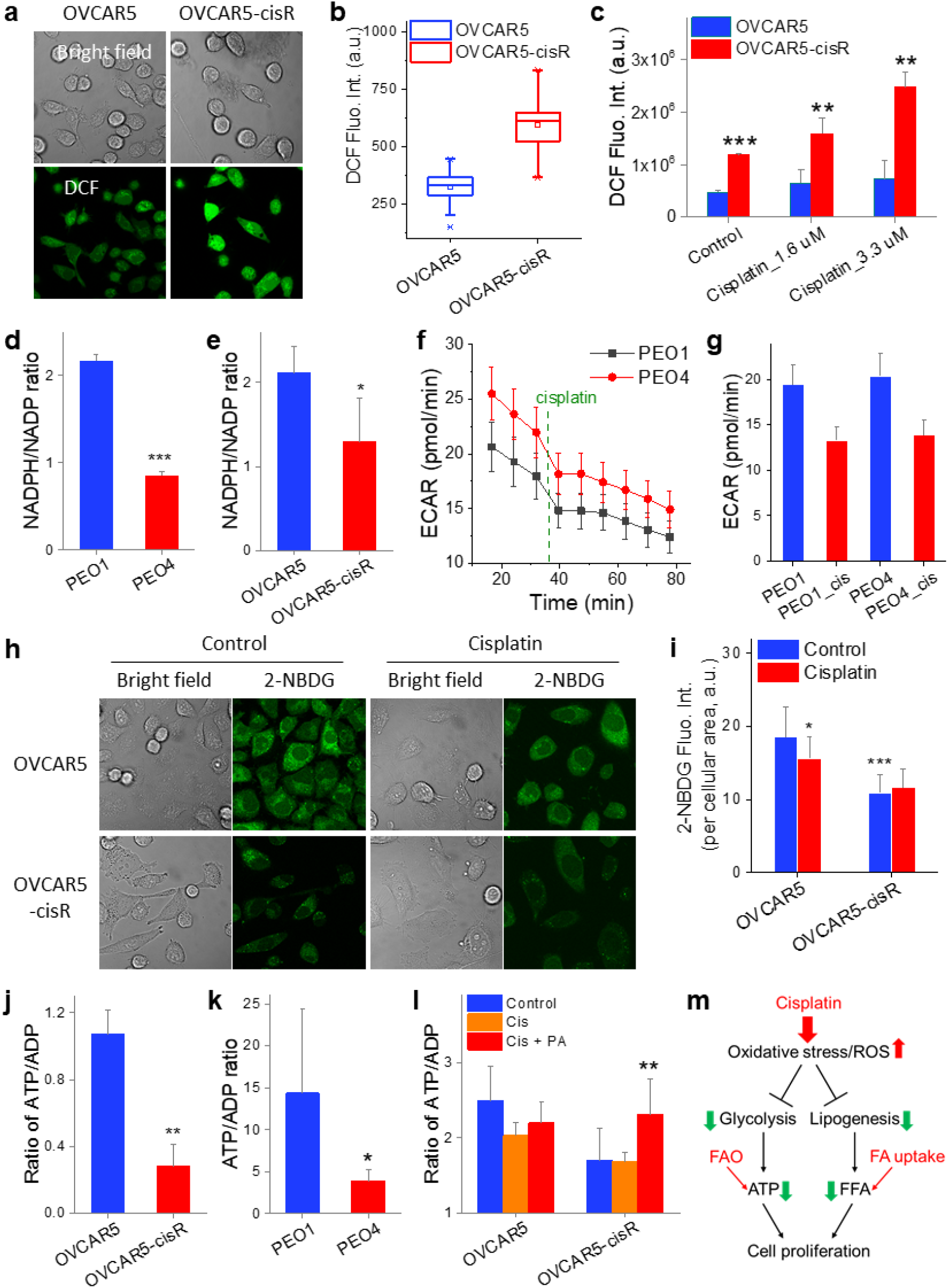
Increased fatty acid uptake and oxidation supports cancer cell survival under cisplatin-induced oxidative stress. **(a)** Representative bright field and fluorescent images of OVCAR5 and OVCAR5-cisR cells after treated with DCFDA cellular ROS assay kit. **(b)** Quantification of DCF fluorescent signal intensity of OVCAR5 and -cisR cells. The outer box indicates 25% to 75% of data; inner box indicates mean; lines represent medium; stars indicate 99% and 1% of data. n ≥ 6. **(c)** Quantification of DCF fluorescent signal intensity of OVCAR5 and -cisR cells with cisplatin treatment at 1.6 μM or 3.3 μM for 24 hours. N ≥ 6. **(d-e)** Quantified NADPH/NADP ratio of cisplatin resistant cell lines and their parental cell lines involving PEO1 and PEO4 (d), and OVCAR5 and -cisR (e). **(f)** ECAR profile of PEO1 and PEO4 after 13.2 μM cisplatin treatment measured by Seahorse. N = 6. **(g)** Quantification of PEO1 and PEO4 cells’ ECAR before and 30 minutes after 13.2 μM cisplatin treatment. N = 6. Representative bright field and fluorescent images of OVCAR5 and -cisR cells with 100 μM glucose analog 2-NBDG treatment for 2 hours after incubation with 3.3 μM cisplatin for 24 hours. **(i)** Quantified 2-NBDG fluorescent signal intensity per cellular area of OVCAR5 and -cisR cells. N ≥ 6. **(j-k)** Quantified ATP/ADP ratio of cisplatin resistant cell lines and their parental cell lines involving OVCAR5 and -cisR (j), and PEO1 and PEO4 (k). N=3. **(l)** Quantified ATP/ADP ratio of OVCAR5 and -cisR treated with 3.3 μM cisplatin with or without supplement of 100 μM palmitic acid for 6 hours. N=3. **(m)** Proposed mechanism about cisplatin effect on cellular metabolism and cell proliferation. The results in all the column graphs are shown as means + SD, n = 8-10. * *P* < 0.05, ** *P* < 0.01, and *** *P* < 0.001.

Next, we examined whether the reduced form of intracellular NADPH is depleted in cisplatin-resistant cells. The results show that NADPH/NADP^+^ ratios are significantly lower in cisplatin-resistant PEO4 (**Fig. 6d**) and OVCAR5-cisR cells (**Fig. 6e**), when compared to PEO1 and OVCAR5 cells, respectively. The reduced NADPH level corroborate our SRS images showing decreased *de novo* lipogenesis in cisplatin-resistant cells. We then analyzed changes in glycolysis by measuring extracellular acidification rate (ECAR) by Seahorse. As anticipated, cisplatin treatment promptly lowered the ECAR rate in both PEO1 and PEO4 cells (**Fig. 6f**), reaching significant reduction within ~30 min of treatment (**Fig. 6g**). On the contrary, cisplatin treatment also induced slight increase of OCR in PEO1 and PEO4 cells (**Supplementary Fig. 6c and S6d**). In agreement with the decreased glycolysis, glucose uptake measured by fluorescent glucose analog, 2-deoxy-2-[(7-nitro-2,1,3-benzoxadiazol-4-yl)amino]-D-glucose (2-NBDG), under confocal microscope was reduced by cisplatin treatment in OVCAR5 cells (**Fig. 6h and 6i**). Further, OVCAR5-cisR cells took up much less 2-NBDG than OVCAR5 cells (**Fig. 6h and 6i**), implying a decreased reliance on glucose metabolism in cisplatin-resistant ovarian cancer cells.

With glycolysis suppressed by increased oxidative stress, we suspected that ATP production will be impaired in cisplatin-resistant cells or cisplatin treated cells. We measured the cellular ATP/ADP level and found the ratio of ATP/ADP was significantly lower in both PEO4 (**Fig. 6j**) and OVCAR5-cisR cells (**Fig. 6k**), when compared to PEO1 and OVCAR5 cells, respectively. Furthermore, acute treatment with cisplatin reduced ATP/ADP ratio in OVCAR5 cells, but not in OVCAR5-cisR cells. Supplementation with palmitic acid significantly increased the ATP level in OVCAR5-cisR cells, but not in OVCAR5 cells (**Fig. 6l**). Collectively, these data reveal metabolic reprogramming of cisplatin-resistant ovarian cancer cells. In summary (**Fig. 6m**), glycolysis and lipogenesis are inhibited by cisplatin-induced oxidative stress, which limits the production of energy as well as synthesis of free fatty acids. To survive and proliferate under cisplatin-induced oxidative stress, cancer cells elevate fatty acid uptake and oxidation as an alternative route of energy production.

### Cisplatin treatment induces a transient metabolic shift toward increased fatty acid uptake in multiple types of cancers

Understanding that increased fatty acid uptake in cisplatin-resistant ovarian cancer cells is likely a stable metabolic reprogramming to adapt to cisplatin induced oxidative stress, we next tested if the same metabolic shift occurs in other types of cancers upon cisplatin treatment. Platinum is widely used across malignancies. Therefore, we selected a few representative cancer cell lines, including MIA PaCa-2 pancreatic cancer, A549 lung cancer and MD-MBA231 breast cancer, to test whether an acute cisplatin treatment changes the rate of fatty acid uptake. We determined the IC50 to cisplatin in these three cell lines and selected 6.6 µM as the final treatment concentration, at which dose no significant cell death was induced (**Supplementary Fig. 7a-c**). Results show that treatment with 6.6 µM cisplatin significantly increased uptake of PA-d_31_ and OA-d_34_ in MIA PaCa-2 cells (**Fig. 7a and 7b**). It is also worth noting that the fold increase in PA uptake was more significant than OA (**Supplementary Fig. 7d**), suggesting PA might be a preferred source of fatty acids for cells under cisplatin induced oxidative stress. Similarly, we observed that treatment with cisplatin also induced a significant increase in PA-d_31_ and OA-d_34_ uptake in A549 (**Fig. 7c-d and S7e**) and MD-MBA231 cells (**Fig. 7e-f and S7f**). These data suggest that our findings are broadly applicable to multiple types of cisplatin-resistant cancers.

**Fig. 7:**
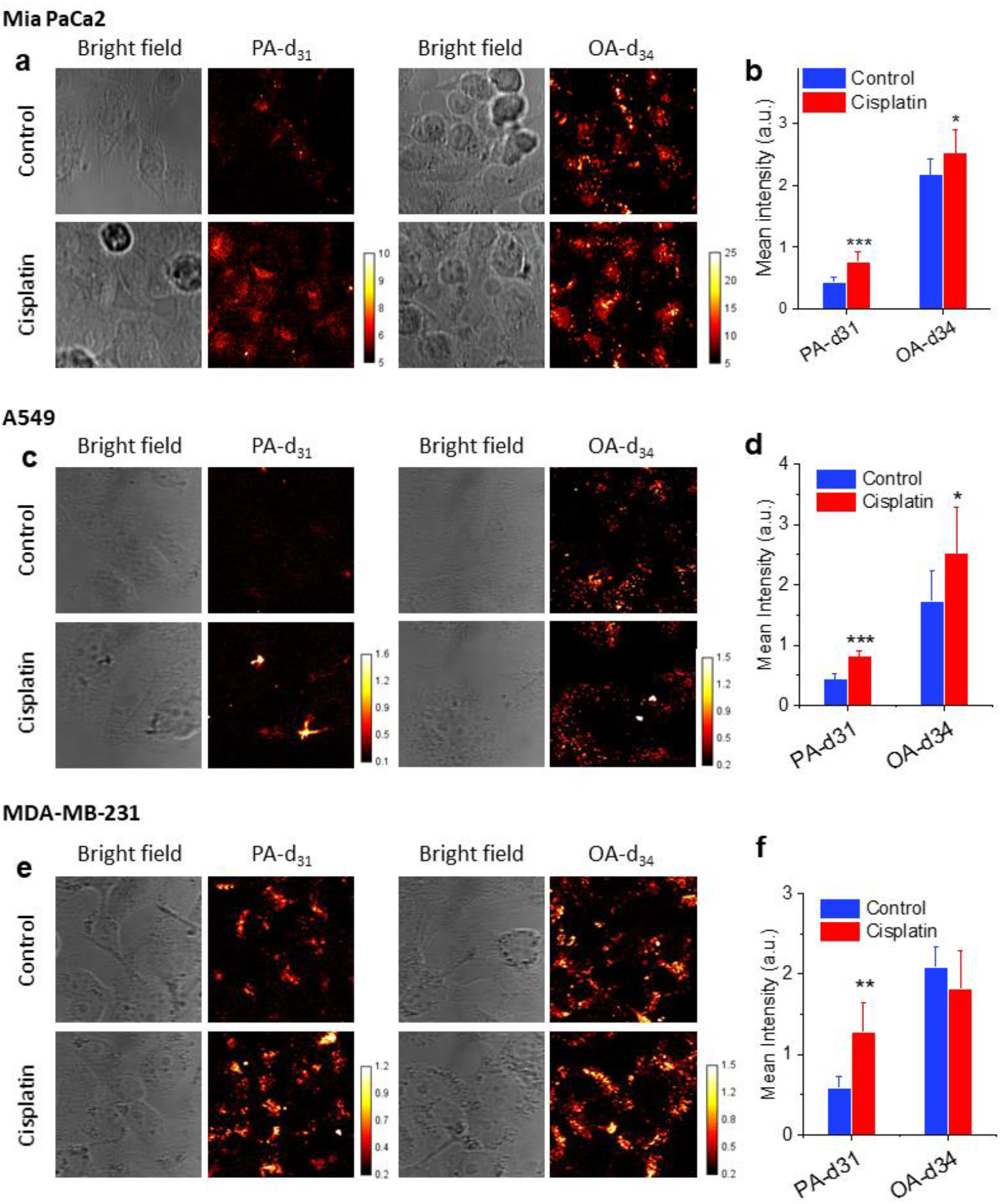
Cisplatin induced fatty acid uptake is a universal metabolic feature in multiple types of cancers. **(a)** Representative bright field and SRS images of Mia Paca2 cells treated with 6.6 μM cisplatin for 24 hours followed by 100 μM PA-d_31_ or OA-d_34_ incubation for 6 h. **(b)** Quantitation of C-D signal in Mia Paca-2 cells treated with or without cisplatin by mean intensity. **(c)** Representative bright field and SRS images of A549 cells treated with 13.2 μM cisplatin for 48 hours followed by 100 μM PA-d_31_ or OA-d_34_ incubation for 6 h. **(d)** Quantitation of C-D signal in A549 cells treated with or without cisplatin by mean intensity. **(e)** Representative bright field and SRS images of MDA-MB-231 cells treated with 6.6 μM cisplatin for 24 hours followed by 100 μM PA-d_31_ or OA-d_34_ incubation for 6 h. **(f)** Quantitation of C-D signal in MDA-MB-231 cells treated with or without cisplatin by mean intensity. The data in all column graphs are shown as means + SD; n ≥ 6. * P<0.05. ** P<0.01. *** P<0.001.

## DISCUSSION

Cisplatin and other platinum-based drugs, such as carboplatin and oxaliplatin, are widely used chemotherapy agents for multiple types of cancers, including ovarian, testicular, bladder, head and neck, non-small-cell lung cancer and others ^53^. Despite high response rate following initial treatment, the effects of cisplatin and carboplatin are limited by severe side effects and high probability of drug resistance development ^53, 54^. In view of the high prevalence of cisplatin or carboplatin use in cancer management, there is a critical need to develop effective therapeutic options to overcome the resistance.

The major mechanism-of-action of cisplatin is formation of DNA-adducts, which block transcription and DNA synthesis, while at the same time, activate DNA damage response mechanisms and mitochondrial detoxification mechanisms. Apoptosis eventually ensues if DNA lesions are not repaired and oxidative stress is not buffered. Numerous efforts have been devoted to elucidating the mechanisms of cancer cell resistance to cisplatin. Most of these studies focused on adduct formation and subsequent activation of cell death pathways, for example, reduced formation of DNA-adducts due to altered uptake/efflux, enhanced DNA damage repair, or impaired mitochondrial apoptosis pathway after adduct formation ^54^. Other mechanisms of cisplatin resistance have received much less attention. Several studies have shown that cisplatin can have another mechanism-of-action by inducing oxidative stress in ovarian ^48, 55^, prostate ^56^, and lung cancer ^57^. Yet, little is known regarding how the cancer cell defending against oxidative stress contributes to cisplatin resistance.

A few recent studies highlighted an association between metabolic reprogramming and cisplatin resistance. An alteration in glycolysis pathway was associated with cisplatin generated oxidative stress in head and neck squamous cell carcinoma ^58^. Lipid droplet production mediated by lysophosphatidylcholine acyltransferase 2 is linked to resistance to oxaliplatin in colorectal cancer ^24^. More recently, adipocyte induced FABP4 was found to mediate carboplatin resistance in ovarian cancer ^59^. In this study using a high-throughput hyperspectral SRS imaging platform, we identified a metabolic switch from glucose to fatty acid dependent anabolic and energy metabolism in cisplatin resistant cancer cells to adapt to cisplatin-induced oxidative stress. As depicted in **Supplementary Fig. 8**, our study reveals decreased glucose uptake, glycolysis and de novo lipogenesis while increased fatty acid uptake and oxidation, suggesting a central metabolic switch in anabolic and energetic metabolism in cisplatin resistant cancer cells. Inhibition of beta-oxidation re-sensitizes cisplatin-resistant ovarian cancer cells to cisplatin treatment both *in vitro* and *in vivo*, paving the foundation towards a new combinational therapy of fatty acid oxidation inhibitors and platinum drugs. We further show that cisplatin treatment induces similar metabolic shift in multiple other types of cancer, implying broad applicability of this strategy.

Fatty acid oxidation, as an alternative path for energy production, has been shown to be upregulated in certain conditions, such as under metabolic stress ^60^. However, it is not known whether and how fatty acid oxidation is deregulated in drug resistant cancer cells. Our data support that cisplatin induced oxidative stress is one of the major mechanisms leading to increased fatty acid uptake and oxidation. Besides fatty acid oxidation, the process of fatty acid uptake could represent another potential target for overcoming cisplatin resistance. However, the regulation of fatty acid uptake involves multiple and redundant transporters, binding proteins and carrier proteins ^41, 42, 44, 61^. Future work is needed to completely elucidate the specific target that mediates the increased fatty acid uptake in cisplatin-resistant cancer cells.

Aside from revealing a potential therapeutic strategy for cisplatin-resistant cancers, quantitative metabolic imaging of *de novo* lipogenesis and fatty acid uptake holds the promise for detection of cisplatin resistance in clinical samples at the single cell level. Based on our study, increased fatty acid uptake accompanied by decreased glycolysis and *de novo* lipogenesis can be a new metabolic marker for detecting cisplatin resistance *ex vivo* in clinical samples. By integrating *de novo* lipogenesis imaging and fatty acid uptake imaging, hyperspectral SRS microscopy can measure a metabolic index, the value of which correlates near linearly to the level of cisplatin resistance in ovarian cancer cells. This finding is confirmed in primary ovarian cancer cells isolated from patients, implicating its potential use in clinical settings. The currently used methods to determine cisplatin resistance rely on cell viability assays or measurement of protein marker expression. These methods could be either time-consuming due to difficulty in culturing primary cells for several generations or inaccurate due to unreliable correlation between protein markers and drug resistance. The metabolic imaging approach reported here provides a fast and quantitative way to determine cisplatin resistance based on functional metabolic signatures in cisplatin-resistant cells.

## Method

### Cell lines

SKOV3 cells were purchased from the American Type Culture Collection (ATCC) and PEO1 and PEO4 were from Sigma Aldrich. OVCAR5 cells were a generous gift from Dr. Marcus Peter, Northwestern University, and COV362 cells were from Dr. Kenneth Nephew, Indiana University. All cell lines were authenticated and tested to be mycoplasma negative. The resistant cell lines SKOV3-cisR, COV362-cisR and OVCAR5-cisR were generated by treatment with 3 or 4 repeated or increasing doses of cisplatin for 24 hours. Surviving cells were allowed to recover for 3 to 4 weeks before receiving the next treatment. Changes in resistance to platinum were estimated by calculating half maximal inhibitory concentration (IC_50_) values as described below ^39^. PEO1, PEO4, OVCAR5, and OVCAR5-cisR cells were cultured in RPMI 1640 medium supplemented with 2 mM L-glutamine, 10% FBS and 100 units/mL penicillin/streptomycin. SKOV3, SKOV3-cisR, COV362 and COV362-cisR cells were cultured in high-glucose DMEM medium supplemented with 10% FBS and 100 units/mL penicillin/streptomycin. For CPT1a knockdown cell lines development, cells were transfected with CPT1a or control shRNA lentiviral transduction particles (Sigma Aldrich) for 48 hours and selected by 1μg/ml puromycin for one week. All cells were cultured at 37°C in a humidified incubator with 5% CO_2_ supply.

## Materials

Glucose-d_7_, palmitic acid-d_31_ (PA-d_31_) and oleic acid-d34 (OA-d_34_) were purchased from Cambridge Isotope Laboratory. 17-Octadecynoic Acid (ODYA), BMS309403, cisplatin, and etomoxir were purchased from Cayman Chemicals. For treatment with cisplatin, 3.3 µM was used as the final concentration, unless otherwise specified.

### Primary human cells

De-identified high grade serous ovarian tumors (HGSOC) and malignant ascites fluid specimens from ovarian cancer patients (n = 5) were obtained at the Northwestern University School of Medicine under an IRB approved protocol (STU00202468). Tumor tissues were enzymatically disassociated into single cell suspensions and cultured as previously described ^62, 63^. After centrifugation at 1,200 rpm for 5 min, 25,000 ascites derived tumor cells were cultured as monolayers in DMEM medium supplemented with 10% FBS and antibiotics prior to SRS imaging.

### PDX *In vivo* model study

Animal studies were approved by the Institutional Animal Care and Use Committee (IACUC) at Northwestern University and were performed in the Developmental Therapeutics Core (DTC) of the Lurie Cancer Center, as previously described ^64^. The platinum-resistant PDX model was previously described. After passage through a donor animal, fresh tumor (equal size) was implanted subcutaneously (SC) in 20 female 7–8-week-old NSG mice (Jackson Labs). Tumor sizes were measured using calipers twice per week and tumor volumes were calculated according to the formula length x width^2^ /2. When the tumor volume reached 100 mm^3^, the animals were randomized into four groups: vehicle, carboplatin alone (10 mg/kg weekly i.p. injection), etomoxir alone (40 mg/kg daily i.p. injection), and combination of carboplatin (10 mg/kg weekly i.p. injection) and etomoxir (40 mg/kg daily i.p. injection). Body weights and habitus were monitored twice per week and mice were sacrificed when largest tumor exceeded 1500 mm^3^ or if human endpoints were reached earlier.

### Large-area hyperspectral stimulated Raman scattering (SRS) imaging

Hyperspectral SRS imaging was performed on a lab-built system following previously published method ^8^. The laser source is a femtosecond laser (InSight DeepSee, Spectra-Physics, Santa Clara, CA, USA) operating at 80 MHz with two synchronized output beams, a tunable pump beam ranging from 680 nm to 1300 nm and a Stokes beam fixed at 1040 nm. For imaging at the C-H vibration region (2800 ~ 3050 cm^−1^), pump beam was tuned to 798 nm. The Stokes beam was modulated at 2.3 MHz by an acousto-optic modulator (1205-C, Isomet). After combination, both beams were chirped by two 12.7 cm long SF57 glass rods and then sent to a laser-scanning microscope. The power of pump and Stokes beam before microscope was controlled to be 20 mW and 200 mW, respectively. A 60x water immersion objective (NA = 1.2, UPlanApo/IR, Olympus) was used to focus the light on the sample, and an oil condenser (NA = 1.4, U-AAC, Olympus) was used to collect the signal. For hyperspectral SRS imaging, a 50-image stack was acquired at different pump-Stokes temporal delay, which was controlled by tuning the optical path difference between pump and Stokes beam through a translation delay stage. Raman shift was calibrated using standard samples, including DMSO, oleic acid, and linolenic acid.

To achieve large-area mapping, samples were fixed on a motorized stage (PH117, Prior Scientific). A lab built LabView based program was used to control moving of the stage and stitching of images. The stage moves to adjacent location with partial overlap after a hyperspectral SRS image was acquired at current location. A montage image composed of 5 × 5 individual 400 × 400-pixel images was acquired at each area of interest. The size of the montage image is approximately 500 × 500 μm. The pixel dwell time is set as 10 μs. For each sample, at least 3 montage images were acquired at different area of interest.

### Spectral phasor and CellProfiler based single cell analysis

The acquired large-area hyperspectral SRS images were segmented through Spectral phasor analysis modified from previously published method ^38^. Spectral phasor was installed as a plugin in ImageJ. The images were transformed into a two-dimension phasor plot based on Fourier Transform. Each dot on the phasor plot represents an SRS spectrum at a particular pixel. Pixels with similar spectra or chemical content were clustered on the phasor plot. “Nuclei” and “lipid” images were generated by mapping the corresponding clusters on the phasor plot back to two separate images.

Lipid analysis in single cells were performed through the software CellProfiler ^65^. The nuclei map and cell image were input into CellProfiler to outline each individual cells. Meanwhile, the lipid map was input into CellProfiler to pick up the lipid droplet (LD) particles. Then, the lipid map was masked onto the outlined cell map to label the lipids by cells. Morphological information of each cell and lipid analysis, including LD number and intensity in each single cell were measured and reported in the output results. The total lipid intensity in each cell was plotted as histogram graphs. For each sample, a few hundreds to a thousand of cells were analyzed.

### Isotope labeling and SRS imaging

For labeling with glucose-d_7_, media was replaced with glucose-free DMEM medium (Thermo Fisher Scientific, # 11966025) + 10% FBS + P/S supplemented with 25 mM glucose-d_7_ after seeding the cells in 35 mm glass-bottom dishes overnight. For labeling with fatty acids or analogs, including PA-d_31_, OA-d_34_ and ODYA, fatty acids or analogs were added to the culture media at final concentration of 100 μM and cells were treated for 6 h. For quantitative SRS imaging, cells on glass-bottom dishes were fixed with 10% neutral buffered formalin for 30 min and washed with PBS for 3 times. Hyperspectral SRS imaging was performed to the cells at Raman spectral region from 2100 to 2300 cm^−1^.

### Reactive Oxidative Species (ROS) measurement

Cellular ROS was measured using a fluorescent probe, 2’,7’-Dichlorofluorescin diacetate (DCFDA) (Sigma Aldrich). Cells seeded in glass-bottom dishes were treated with or without 3.3 µM cisplatin for 3h. DCFDA was added to the medium at final concentration of 10 µM and incubated for 15 min. After washing with PBS for 3 times, cells were immediately imaged under confocal microscope (Zeiss LSM 700 microscope) with 488 nm as the exciting source. Laser power was controlled at low setting to avoid fluorescence extinction. Images at ~10 field of view were acquired for each sample.

### Fatty acid oxidation assay

Fatty acid oxidation was measured using a commercial kit (Abcam, #ab217602) following the provided protocol. Briefly, cells were seeded in 96-well plate with 150k cells/well. After incubation overnight, medium was replaced by freshly prepared medium. 10 μL of Extracellular O_2_ consumption reagent, and 2 drops of high-sensitivity mineral oil (pre-heated at 37 °C) were added to each well. Fluorescence was measured in plate reader at 2 min intervals for 180 min at excitation/emission = 380/650 nm. Etomoxir was added at final concentration of 40 μM to block the fatty acid oxidation. Oxygen consumption rate (OCR) was presented as Δ_Fluorescence intensity_/min/cell and fatty acid oxidation rate is calculated as OCR_FAO_ = OCR_total_ – OCR_Etomoxir_. At least three replicates were included for each measurement.

### NADPH and ATP assay

NADP/NADPH and ADP/ATP were measured using commercial kits (Abcam, #ab65349 and #ab65313). For NADPH measurement, cells (~1×10^6^ cells) were pelleted and extracted using NADPH/NADP extraction buffer. Centrifuge at 10,000 g for 10 min and collect the supernatant. Total NADP/NADPH was directly measured using the assay kit and NADPH alone was measured by firstly decomposing NADP by heating at 60°C for 30 min. Absorption at 450 nm was measured by plate reader at 2h after development. For ATP measurement, cells were seeded in 96-well plates. ATP was measured directly and total ATP + ADP was measured by converting ADP to ATP. Luminescent signal was measured by plate reader 2 min after mixed. At least three replicates were included for each measurement.

### Cell viability assay

Cell viability was measured by MTS assay (Abcam, #ab197010) or by CellTiter-Glo assay (Promega, #G7570). Cells were seeded at 96-well plates at densities of 2000~5000 cells per well overnight. Treatment was added to the cells at indicated concentrations for 72 h. Cell viability was measured by incubating with MTS reagent for 4 h and reading absorbance at 490 nm or incubating with CellTier-Glo reagent for 10 min and reading luminescence by a plate reader. Six replicates were used for each group.

### Glucose uptake assay

Glucose uptake was measured using a fluorescent glucose analog, (2-NBDG) (Cayman chemicals). Cells seeded in glass-bottom dishes were incubated with 100 µM 2-NBDG for 2h. Fluorescent images were taken by confocal microscope (Zeiss LSM 700 microscope) with 488 nm laser as excitation source. Images at ~10 field of view were acquired for each sample.

### Measurement of OCR and ECAR by Seahorse

Cell lines were seeded in a Seahorse XF96 Cell Culture Microplate (Agilent) at density of 2×10^4^ (OVCAR5 pair) or 4×10^4^ (PEO pair) per well. After incubation at 37 °C overnight for cell attachment, oxygen concentration rate (OCR) and extracellular acidification rate (ECAR) were measured through Seahorse XFe96 Analyzer (Agilent). Measurement time was 30 seconds following 3 minutes mixture and 30 seconds waiting time. First three cycles were used for basal respiration measurement. 25 μl treatment in load cartridge was injected to the well after first three cycles. 26.4 μM cisplatin and 40 μM etomoxir treatment’s effect on OC’s OCR and ECAR have been investigated.

### Reverse transcription-PCR (RT-PCR)

Total RNA from ovarian cancer cell lines were extracted via RNeasy Mini Kit (Qiagen Inc.) and reverse transcribed by iScript cDNA Synthesis Kit (Bio-Rad). RT-PCR was performed through StepOne Plus RT-PCR (Applied Biosystems) using Power SYBR Green Master Mix (Thermo Fisher Scientific) All procedure was following manufactures’ instructions. RT-PCR reaction generated a melting curve and cycle threshold (Ct) was recorded for the gene of interested and house-keeping control gene (PPIA). The relative RNA expression level was calculated as ΔCt and normalized by subtracting the Ct value of target gene from that of control gene. Results are presented as means + SD. Measurements were performed in biological triplicate and each biological replicate includes three technical replicates.

### Western blot

Proteins were extracted from cell culture by RIPA lysis buffer (Sigma Aldrich) with protease and phosphatase inhibitor cocktail and sample reducing agent (Thermo Fisher Scientific). Proteins were separated in Bolt Bis-Tris Plus gels (Thermo Fisher Scientific) through gel electrophoresis and transferred to PVDF membrane (Bio-Rad). After blocking in 5% non-fat milk (Bio-Rad) for 1 hour at room temperature, membranes were incubated with primary antibodies (CPT1a (1:1000) (Proteintech) and GapDH (1:2000) (Cell Signaling Technology)) overnight at 4°C followed by secondary anti-mouse antibodies (1:10000) (Cell Signaling Technology) for 1 hour at room temperature. Protein bands were developed by ECL reagent (Thermo Fisher Scientific) and detected through ChemiDoc MP imaging system (Bio-Rad). The band intensity was determined using ImageJ.

### Quantification and Statistical Analysis

All the data are presented as means ± SD unless otherwise specified. The statistical significance was analyzed using Student’s t test with two-tailed. All experiments were repeated at least 3 times. N is indicated sample size for each experiment. *P*<0.05 was considered statistically different. Statistical parameters can be found in thefigure legends.

## Acknowledgements

This work was supported by R01CA224275 to DM and JXC and R33CA223581 and R35GM136223 to JXC. Research reported in this publication was supported by the Boston University Micro and Nano Imaging Facility and the Office of the Director, National Institutes of Health of the National Institutes of Health under award Number S10OD024993. PDX experiments were performed in the Northwestern University –Center for Developmental Therapeutics supported by the Cancer Center Support Grant NCI CA060553.

## Author Contributions

J.L., J.X.C. and D.M. co-designed the experiments. J.L., Y.T., H.C., G.Z. performed the experiments. K.C.H. provided help in data analysis. J.L. and Y.T. wrote the manuscript. All authors read and edited the manuscript.

## Additional Information

Supplementary is available for this paper.

## Correspondence and requests for materials

should be addressed to J.X.C or D.M.

