## Supplementary Information for "Metabolic Reprogramming from Glycolysis to Fatty Acid Uptake and beta-Oxidation in Platinum-Resistant Cancer Cells"

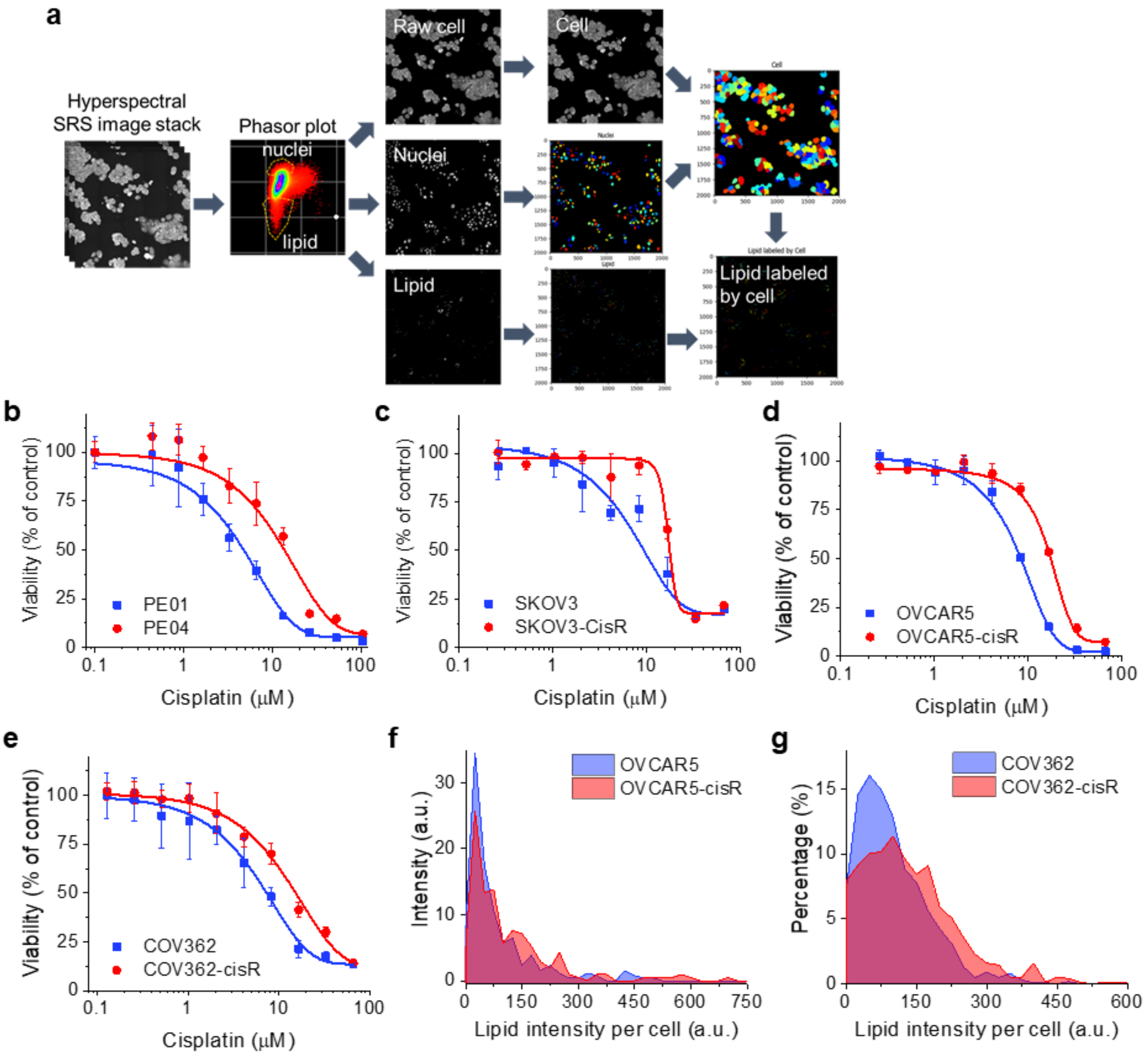


**Supplementary Fig. 1. High-throughput profiling of lipid metabolism in ovarian cancer cell lines. (a)** Image processing flow for high-throughput single cell analysis of lipids. A hyperspectral SRS image stack was segmented through spectral phasor analysis to generate segmented images of nuclei and lipid, which together with raw cell image were input in CellProfiler to outline each individual cells. Lipids were color-coded based on the colors of their parental cells. Quantitative analysis of cell morphology, lipid quantity and intensity were also produced along with the images. The image area is 500 µm by 500 µm. **(b-e)** Dose-response to cisplatin in PEO1 & 4 (b), SKOV3 & -cisR (c), OVCAR5 & -cisR (d), COV362 & -cisR cells (e). The results are shown as means±SD, n = 6. **(f-g)** Histograms of integrated cellular lipid intensity in OVCAR5 & -cisR (f) and COV362 & -cisR cells (g).

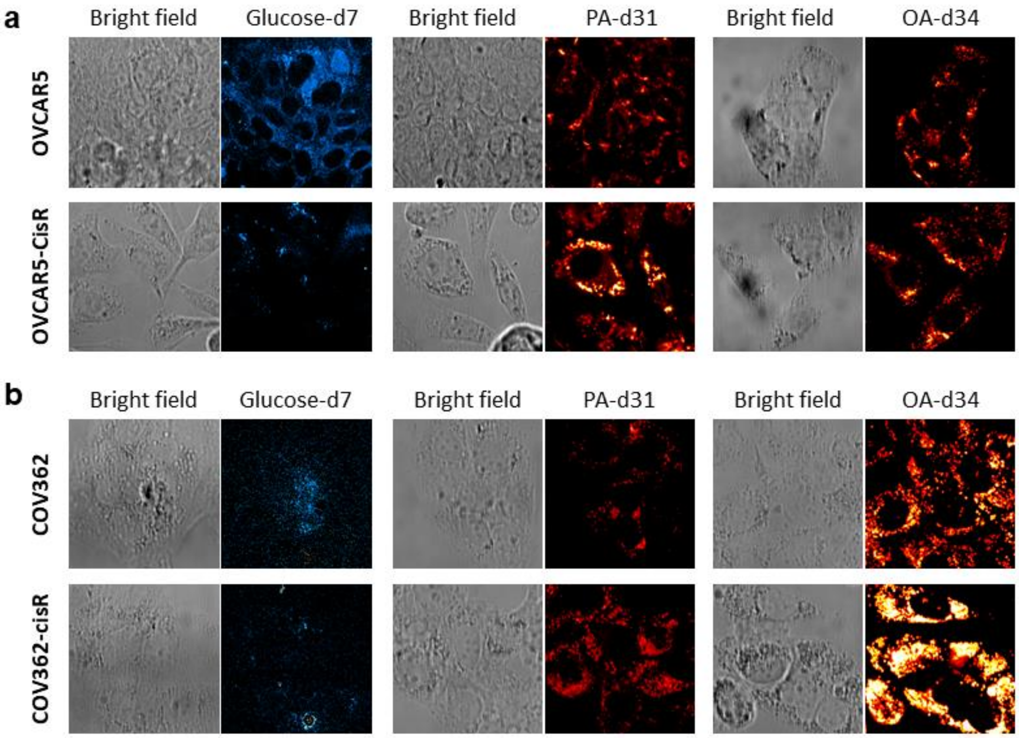


**Supplementary Fig. 2. Increased fatty acid uptake, not de novo lipogenesis, is the major contributor to lipid accumulation in cisplatin-resistant ovarian cancer cells. (a)** Representative bright field and SRS images of OVCAR5 and OVCAR5-cisR cells fed with glucose-$d_7$ for 3 days, PA-$d_{31}$ for 6 h, or OA-$d_{34}$ for 6 h. **(b)** Representative bright field and SRS images of COV362 and COV362-cisR cells fed with glucose-$d_7$ for 3 days, PA-$d_{31}$ for 6 h, or OA-$d_{34}$ for 6 h.

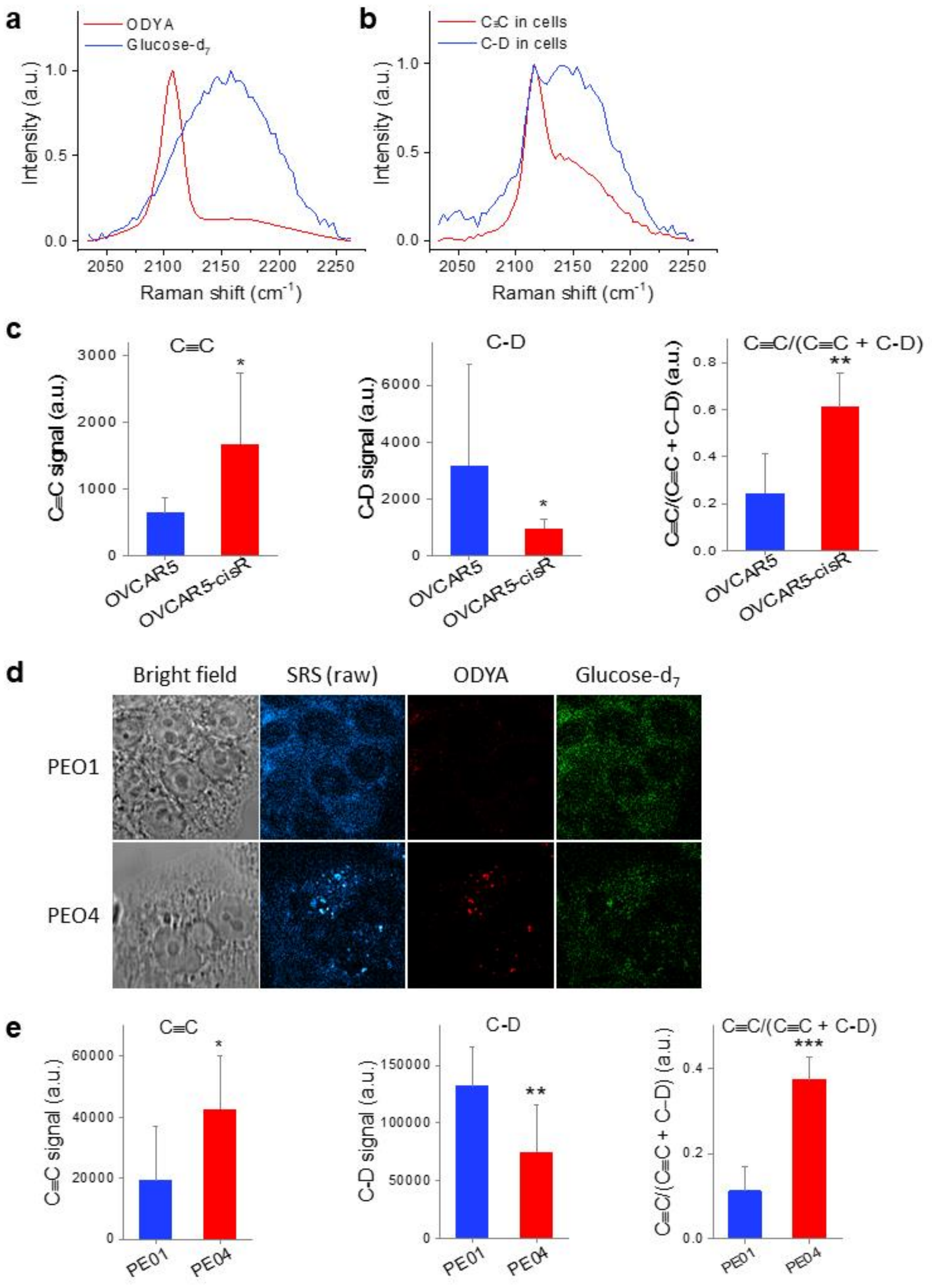


**Supplementary Fig. 3. Metabolic index by integrating glucose derived lipogenesis and fatty acid uptake directly correlates with cisplatin resistance. (a)** Normalized SRS spectra of ODYA and glucose-d7 in cells. **(b)** Output SRS spectra from phasor analysis of C≡C bonds from ODYA and C-D bonds from glucose-$d_7$ and metabolites. **(c)** Quantitative analysis of ODYA derived C≡C intensity, glucose-$d_7$ derived C-D intensity, and the ratio of C≡C/(C≡C + C-D) in OVCAR5 and OVCAR5-cisR cells. **(d)** Representative bright field images, raw SRS images, and processed SRS images of ODYA and glucose-$d_7$ in PEO1 and PEO4 cells. **(e)** Quantitative analysis of ODYA derived C≡C intensity, glucose-$d_7$ derived C-D intensity, and the ratio of C≡C/(C≡C + C-D) in PEO1 and PEO4 cells. The results in all the column graphs are shown as means + SD, n = 8-10. * $P < 0.05$, ** $P < 0.01$, and *** $P < 0.001$.

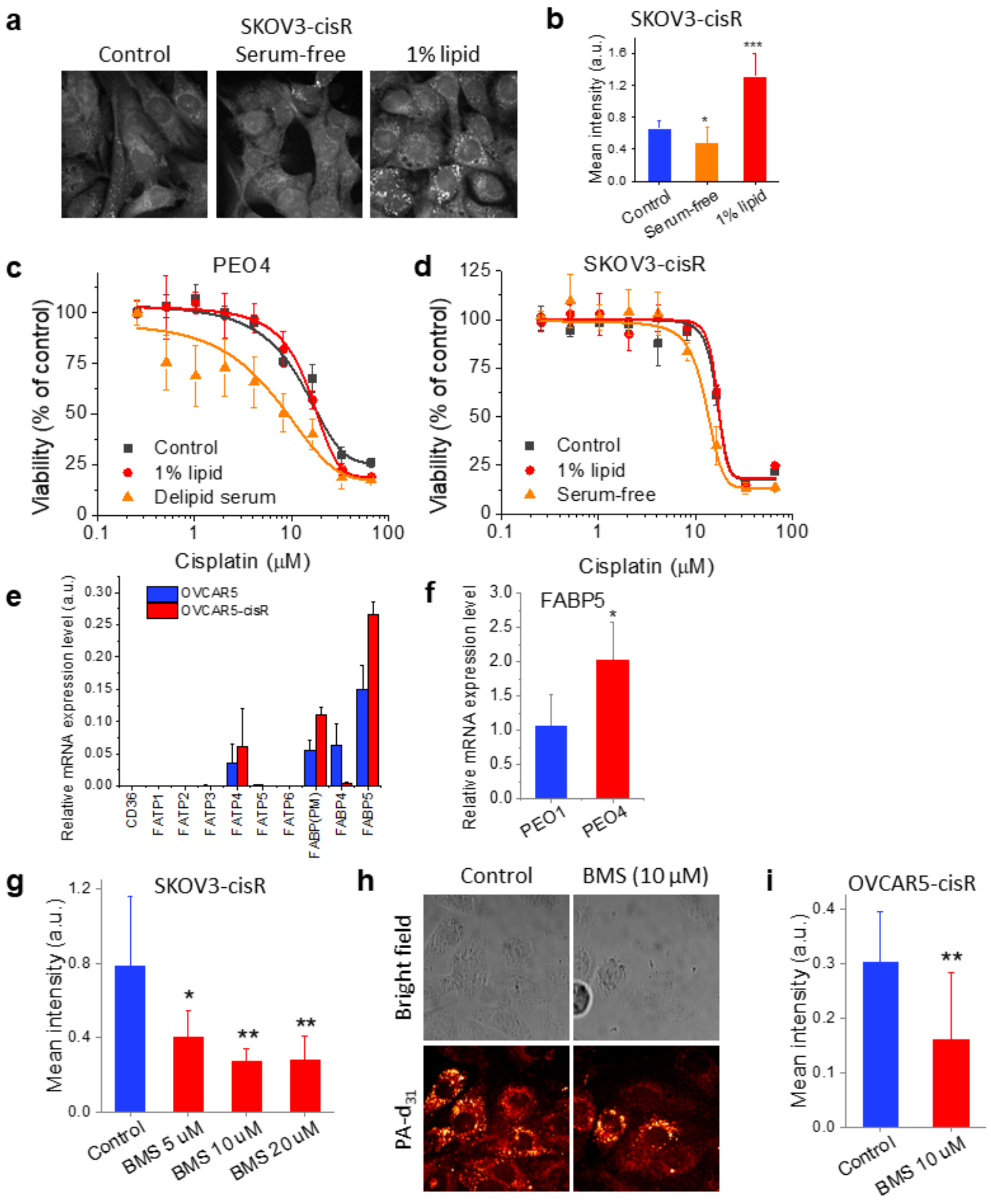


**Supplementary Fig. 4. Fatty acid uptake directly contributes to cisplatin resistance. (a)** Representative SRS images of SKOV3-cisR cell cultured with control serum (FBS), no serum and control serum supplemented with 1% lipid mixture. **(b)** Quantitative C-H signal from lipid droplet of SKOV3-cisR cell cultured with control serum (FBS), no serum or control serum supplemented with 1% lipid for 24 hours. **(c-d)** Dose-response to cisplatin under culture environment with control, reduced (medium containing delipid serum or no serum) and increased (control serum supplemented with 1% lipid mixture) lipid content for PEO1 (c), SKOV3 (d). The data are shown as means ± SD; n = 3. **(e)** Relative mRNA express level of CD36, FATP1-6, FABP4-5, and GOT2 (FABP PM) in OVCAR5 and -cisR cells. The data are shown as means + SD; n = 3. **(f)** Quantification of C-D SRS signal intensity from SKOV3-cisR after BMS treatment at 5 μM, 10 μM or 20 μM for 24 hours with the incubation of 100 μM PA-d31 for 6 hours. **(g)** Representative bright field and SRS images of OVCAR5-cisR cell after BMS treatment at 10 μM for 24 hours with the incubation of 100 μM PA-d31 for 6 hours. **(h)** Quantification of C-D SRS signal intensity from OVCAR5-cisR cell after BMS treatment at 10 μM for 24 hours with the incubation of 100 μM PA-d31 for 6 hours. The results in SRS signal intensity quantification are shown as means + SD; n = 8-10. * $P < 0.05$, ** $P<0.01$.

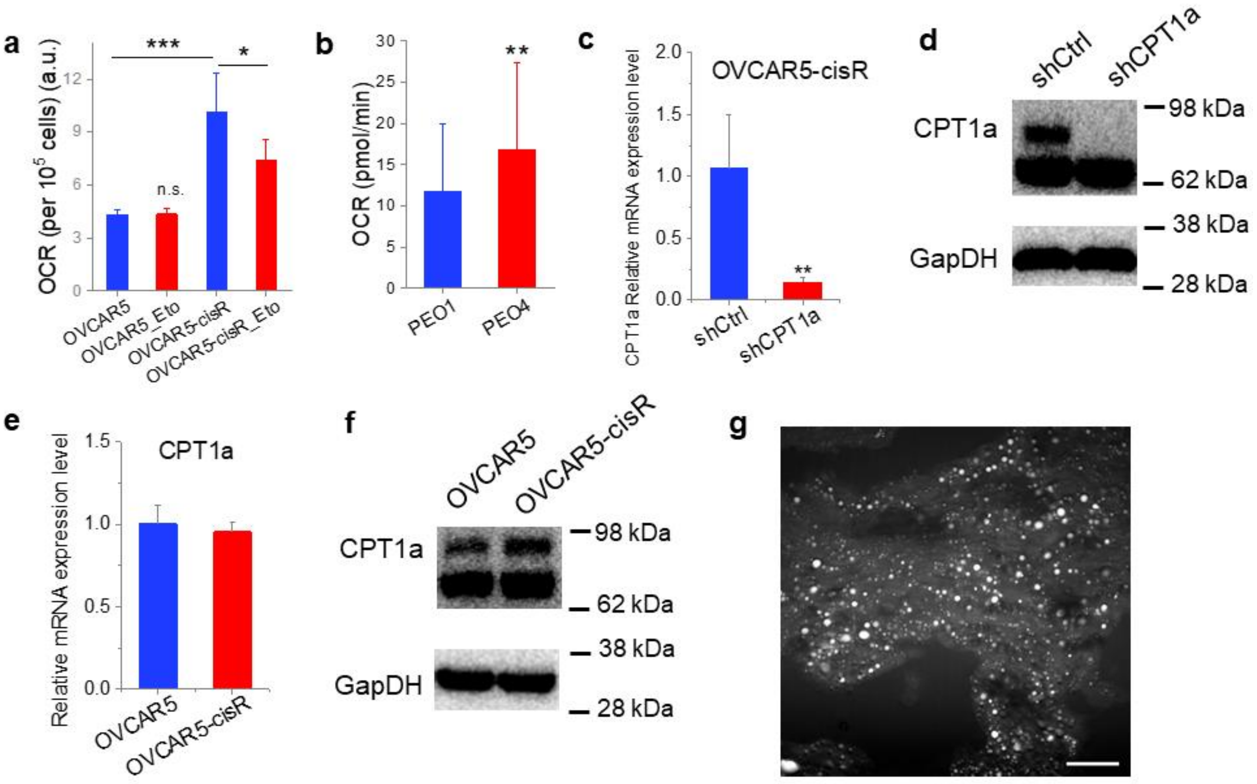


**Supplementary Fig. 5. Fatty acid uptake contributes to cisplatin resistance by increasing fatty acid oxidation. (a)** Quantification of OCR for OVCAR5 and OVCAR5-cisR cells with or without etomoxir treatment at 40 μM measured by extracellular oxygen consumption kit (Abcam). **(b)** Quantification of OCR for PEO1 and PEO4 cells measured through Seahorse XF Analyzers (Seahorse Bioscience). The data are shown as means + SD; n = 6. **(c)** Relative mRNA express level of CPT1a in OVCAR5-cisR shCtrl and shCPT1a cell. **(d)** Western Blot of CPT1a and GapDH for OVCAR5-cisR shCtrl and shCPT1a cell. **(e)** Relative mRNA express level of CPT1a in OVCAR5 and OVCAR5-cisR cell. **(f)** Western Blot of CPT1a and GapDH for OVCAR5 and OVCAR5-cisR cell. The mRNA expression result results are shown as means + SD; n = 3-4. * $P<0.05$. ** $P<0$. *** $P<0.001$. n.s. $P>0.05$. **(g)** Representative SRS image of ovarian tumor tissue slide from PDX model. The scale bar represents 20μm.

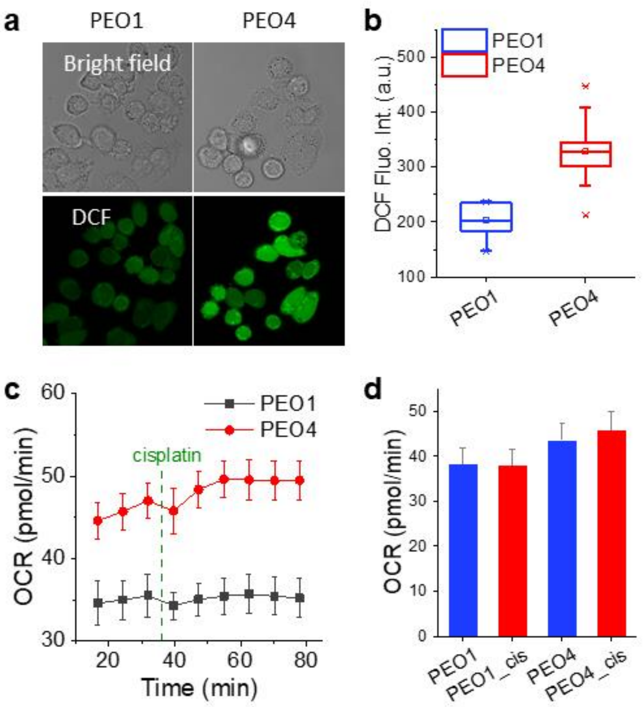


**Supplementary Fig. 6. Increased fatty acid uptake and oxidation supports cancer cell survival under cisplatin-induced oxidative stress. (a)** Representative bright field and fluorescent images of PEO1 and PEO4 cells after treated with DCFDA cellular ROS assay kit. **(b)** Quantification of DCF fluorescent signal intensity for PEO1 and PEO4 cells. The outer box indicates 25% to 75% of data; inner box indicates mean; line represents medium; stars indicate 99% and 1% of data. n ≥ 6. **(c)** OCR profiles of PEO1 and PEO4 after 13.2 μM cisplatin treatment measured by Seahorse. **(d)** Quantification of OCR for PEO1 and PEO4 cells before and 30 minutes after 13.2 μM cisplatin treatment measured by Seahorse. The data in all OCR measurement are shown as means + SD; n = 6.

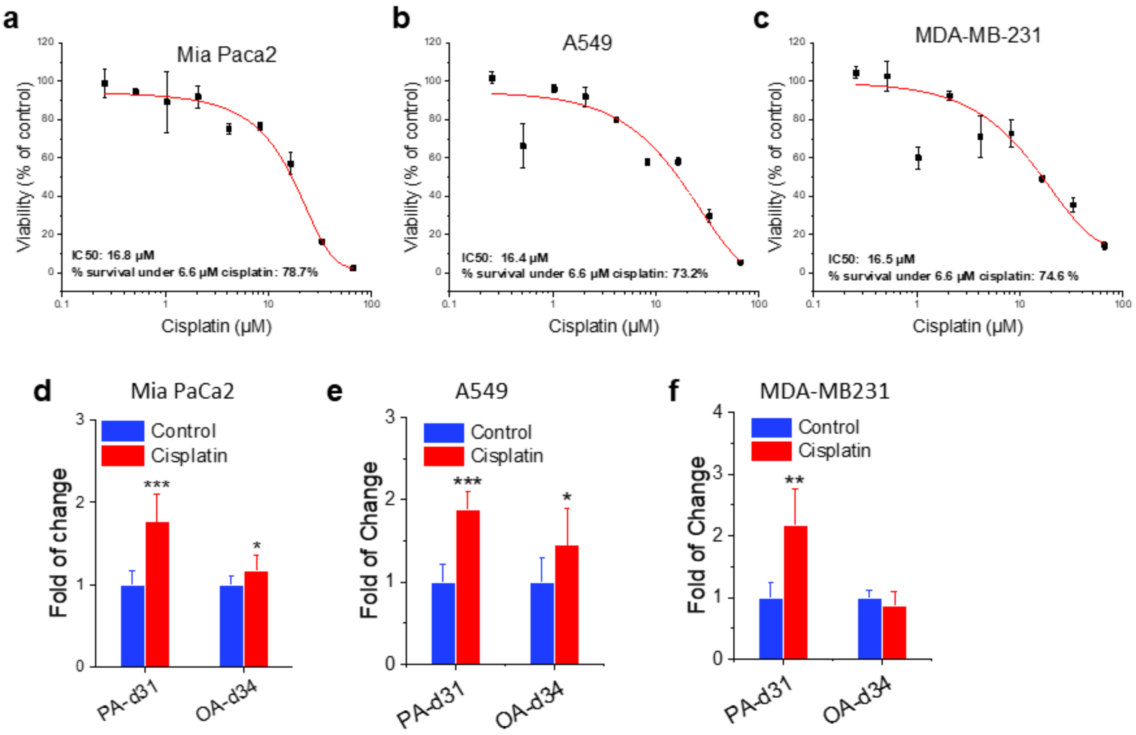


**Supplementary Fig. 7. Cisplatin induced fatty acid uptake is a universal metabolic feature in multiple types of cancers. (a-b)** Dose-response to cisplatin for Mia Paca2 (a) A549 (b) and MD-MBA231 (c). The data are shown as means ± SD; n = 3. **(d-f)** Quantitation of C-D signal in Mia Paca2 (d), A549 (e), and MDA-MB-231 (f) cells treated with or without cisplatin by fold of change. The data in all column graphs are shown as means + SD; n ≥ 6. * P<0.05. ** P<0.01. *** P<0.001.

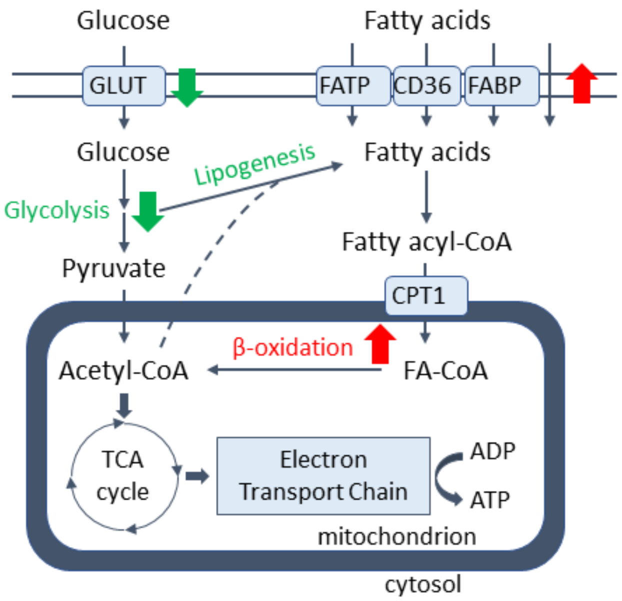


**Supplementary Fig. 8. Diagram showing metabolic reprogramming in cisplatin-resistant ovarian cancer cells.** Proposed mechanism about cellular metabolism switch from glycolysis to fatty acid β-oxidation in cisplatin-resistant ovarian cancer cell.

**Supplementary Table 1. Summary of quantitation results of glucose-$d_7$, PA-$d_{31}$, and OA-$d_{34}$ and $IC_{50}$s of cisplatin in 4 pairs of parental and cisplatin-resistant ovarian cancer cells.**

| Cell lines | $IC_{50}$ of cisplatin (µM) | Intensity (area fraction (%) | | |
|---|---|---|---|---|
| | | G-d7 | PA-d31 | OA-d34 |
| PE01 | 4.72 | 3.88 | 6.11 | 17.25 |
| PE04 | 13.57 | 0.93 | 13.69 | 21.61 |
| SKOV3 | 10.07 | 3.39 | 9.88 | 6.24 |
| SKOV3-CisR | 17.29 | 0.84 | 15.79 | 10.97 |
| OVCAR5 | 8.26 | 3.64 | 10.37 | 18.35 |
| OVCAR5-CisR | 17.43 | 2.51 | 15.91 | 23.28 |
| COV362 | 7.19 | 3.14 | 4.44 | 31.00 |
| COV362-cisR | 15.17 | 2.03 | 14.88 | 41.43 |

Quantitation of glucose-d7, PA-d31, and OA-d34 are shown as area fraction of C-D signal out of total cellular area. Only the mean values are shown.

**Supplementary Table 2. Primer sequences for RT-PCR measurement.**

| Gene name | Forward sequence | Backward sequence |
|---|---|---|
| CPT1a | TCCAGTTGGCTTATCGTGGTG | TCCAGAGTCCGATTGATTTTTGC |
| FABP5 | TGAAGGAGCTAGGAGTGGGAA | TGCACCATCTGTAAAGTTGCAG |
| FABP (PM) | GGAAGGAAATAGCAACAGTGG | TCCTACACGCTCACCATATAAGC |
| FATP1 | CTTCGATGGCTATGTCAGCGA | AGCACGTCACCTGAGAGGTAG |
| FATP2 | ATGCGAGAAAAGTTGGTGCT | TTTCATCACGGACAGGTTCA |
| FATP3 | ATACCTGGGAGCGTTTTGTG | CCGCTGTCCTGTGTAGTTGA |
| FATP4 | CTTTTCCAGCCGCTTCCACA | TGGCTGGCAGGGAATGCA |
| FATP5 | AGCTCCTGCGGTACTTGTGT | AAGGTCTCCCACACATCAGC |
| FATP6 | GCTGGGCCTTATAAGCACACA | CAACCTCAGTGGTTGCGACA |
| CD36 | GGCTGTGACCGGAACTGTG | AGGTCTCCAACTGGCATTAGAA |
| FABP4 | ACTGGGCCAGGAATTTGACG | CTCGTGGAAGTGACGCCTT |
| PPIA | CCCACCGTGTTCTTCGACATT | GGACCCGTATGCTTTAGGATGA |
